# KobA mediates kinesin-1-dependent transport of dynein-dynactin to microtubule plus ends

**DOI:** 10.64898/2026.11.02.763303

**Authors:** Rongde Qiu, Marisa Delgado, Jun Zhang, Mario Pinar, Paula Polonìo, Miguel A. Peñalva, Xin Xiang

## Abstract

Cytoplasmic dynein transports diverse cargoes toward microtubule minus ends following activation by the dynactin complex and cargo adapters. In elongated cells such as fungal hyphae and neurons, dynein must first undergo kinesin-1-dependent transport to microtubule plus ends to engage cargoes, but the mechanism linking these opposing motors remains poorly understood. Here, we identify KobA in the filamentous fungus *Aspergillus nidulans* as a kinesin-1-binding, coiled-coil protein essential for kinesin-1 (KinA)-mediated plus-end transport of dynein and dynactin. KobA binds the C-terminal cargo-binding domain of KinA. Interestingly, KobA is not required for the general function of KinA in supporting hyphal growth alongside myosin-V, but is specifically required for the microtubule plus-end accumulation of dynein-dynactin even when KinA is activated. Mechanistically, KobA enhances the interaction between KinA and dynactin-dynein. Within its disordered N-terminal region, KobA harbors a Spindly motif typically present in dynein-activating cargo adapters that engage the dynactin pointed end. AlphaFold-3 predictions suggest that the Spindly motif of KobA also binds the dynactin pointed end, and in support of this notion, point mutations within the KobA Spindly motif disrupt the KinA–dynactin association, causing dynein to decorate microtubules rather than accumulate at plus ends. Together, our findings establish KobA as a novel adapter that utilizes a classic cargo adapter Spindly motif to link kinesin-1 to dynein-dynactin, ensuring targeted plus-end delivery of the minus-end motor.

## Introduction

The microtubule minus-end-directed motor cytoplasmic dynein transports a variety of cargoes, and dynein activation requires LIS1, NudE and the dynactin complex, as well as specific cargo adapter proteins ^1–4^. Prior to binding cargo adapters that activate the minus-end-directed movement of dynein ^5,6^, cytoplasmic dynein and dynactin can be observed to accumulate at the microtubule plus end in various cell types ranging from fungi to cultured mammalian cells ^7–15^ as well as in vitro reconstituted systems with dynamic microtubules ^1^^6,17^. In cultured neuronal cells, although the plus-end accumulation of dynein and dynactin has not yet been documented, both dynein and dynactin are able to move in a plus-end directed fashion ^18–20^.

The plus-end-directed kinesins have been implicated in delivering dynein to the microtubule plus ends ^21^. In the budding yeast, kinesin-7 (Kip2)-mediated transport of dynein contributes to its accumulation at the microtubule plus ends ^22,23^, although dynein can also be directly recruited to the plus ends from the cytosol ^24^. In filamentous fungi such as *Aspergillus nidulans* and *Ustilago maydis*, kinesin-1s are important for the plus-end accumulation of dynein and dynactin ^14,15,25–28^. The fact that kinesin-1 is critical for dynein-mediated transport of early endosomes away from the tip, where endocytosis predominates, highlights the physiological importance of kinesin-1-mediated dynein localization at the plus ends ^15,26,29,30^.

In higher eukaryotic systems such as *Drosophila melanogaster* oocytes, *Caenorhabditis elegans* neurons and mammalian neurons, kinesin-1 also plays a role in transporting dynein or dynactin toward the microtubule plus ends, where dynein can initiate the retrograde transport of its cargoes ^18,19,31–36^. Kinesin-1 in higher eukaryotic cells transports a variety of cargoes and its activity is regulated by multiple factors including but not limited to cargo adaptors and kinesin light chains ^37–55^. Kinesin-1s in *C. elegans* and mammalian neurons possibly use different mechanisms to get dynein to the plus ends ^18,19,31^. For example, while JIP3, a dynein-activating cargo adapter ^56–59^, is involved in linking dynein and kinesin-1 in *C. elegans* neurons ^31^, the direct interactions of both the heavy and light chains of kinesin-1 with the dynein intermediate chain are implicated in transporting dynein to the plus end in cultured mammalian neurons ^19,60^. In filamentous fungi, kinesin light chains have not been found ^61,62^. In *A. nidulans*, the only kinesin-1 is KinA ^63^, and evidence suggests that the dynactin complex enhances the association between dynein and KinA ^64^. However, exactly how kinesin-1 transports dynein and dynactin in vivo is still not well understood.

In the current study, we identified a kinesin-1/KinA-binding protein in *A. nidulans*, which we named KobA (<u>K</u>inesin-<u>o</u>ne-<u>b</u>inding protein in *<u>A</u>spergillus nidulans*), needed for KinA-dependent dynein localization at the microtubule plus end. In the Δ*kobA* mutant, GFP-labeled dynein and dynactin proteins could no longer accumulate at the microtubule plus ends. However, while KinA is almost essential for colony growth in a strain without myosin-V ^27,65^, due to the overlapping functions of the motors in the transport of secretory vesicles, KobA is dispensable for this function. Thus, KobA is specifically required for kinesin-1-mediated transport of dynein-dynactin toward the microtubule plus ends. Furthermore, we identified a Spindly motif in KobA critical for the plus-end accumulation of dynein. The Spindly motif is normally found in dynein-activating adapters and it interacts with the pointed end of the dynactin complex ^66–69^. We found that KobA helps the connection between KinA and dynein-dynactin, and the Spindly motif of KobA plays an important role in this interaction. Thus, we have identified a kinesin-1-binding protein that interestingly uses a Spindly motif to ensure the plus-end-directed transport of dynein and dynactin.

## Results

### Identification of KobA as a protein that binds the C-terminal cargo-binding domain of kinesin-1 (KinA)

KobA (An2744) in *A. nidulans* was identified as a KinA-associated protein by MS/MS shotgun sequencing of proteins that co-purified with the GFP-labeled cargo-binding domain (CBD, aa731-927) of KinA (around the CC4 region) but not with the globular tail domain (GTD) of myosin V (Figure 1A). Reciprocal pulldowns showed that KinA is pulled down with KobA fused either to GFP or an S-tag (Figure 1A), confirming that this was a meaningful interaction. KobA is a 414 aa protein with an unstructured N terminus (aa1-166), which contains amino acids LADEL (aa77-81) that resemble a “Spindly motif” in dynein adapters known to interact with dynactin pointed-end protein p25 ^57,66–70^ (Figure 1B).

**Figure 1.**
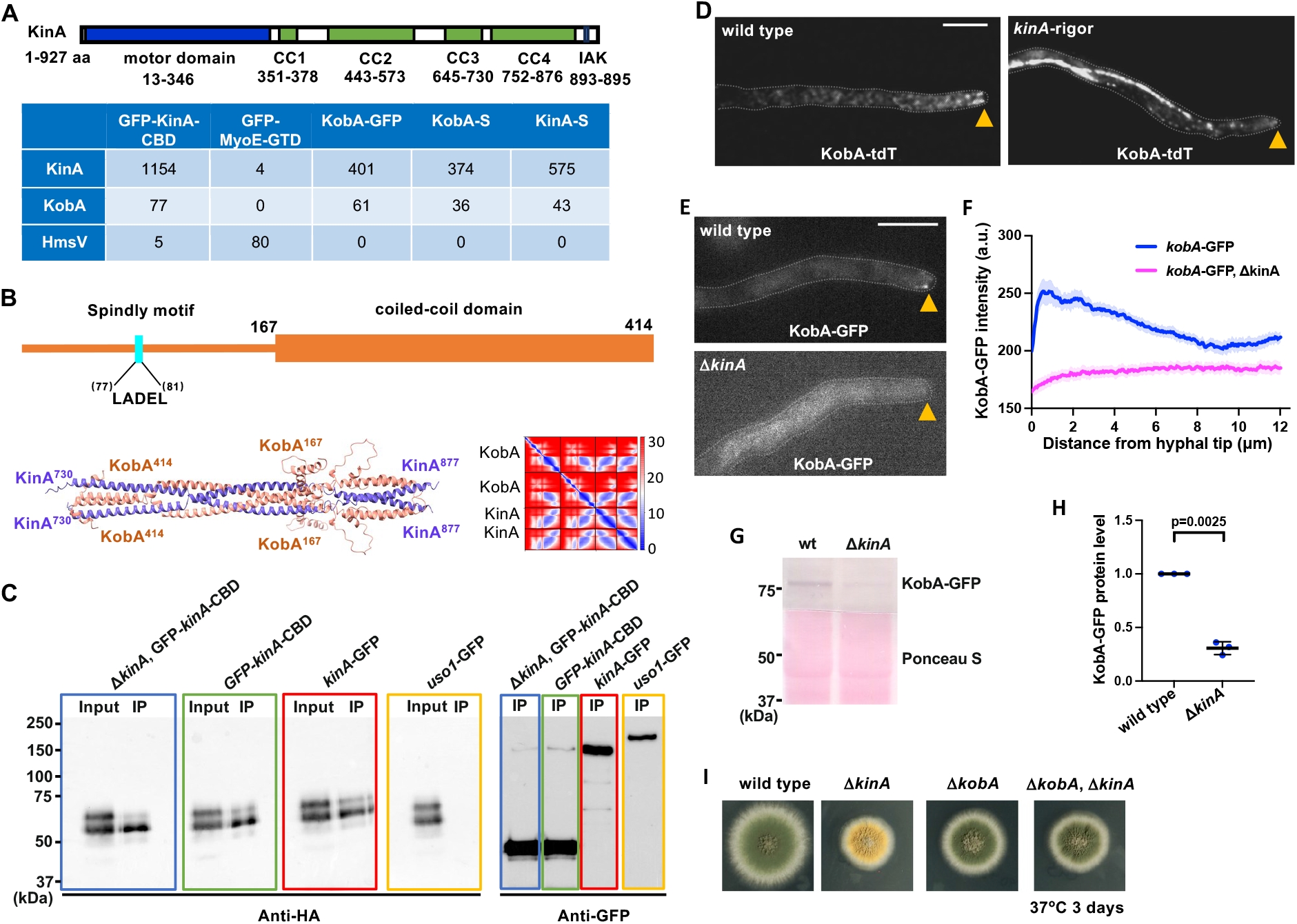
The KobA-KinA association indicated by western blots and AlphaFold-3 prediction. (A) A cartoon of KinA domains and mass spectrometry data showing KinA-KobA association as evidenced by reciprocal pulldowns. Note that the cargo-binding domain (CBD) of KinA used in this study is aa 731-927, which covers the KinA CC4 domain and the C-terminus. Mass spectrometry data showing peptide spectral matches (PSM) of proteins pulled down with GFP-labeled CBD of KinA (GFP-KinA-CBD), GFP-labeled globular tail domain (GTD) of myosin-V (MyoE in *A. nidulans*) (GFP-MyoE-GTD) ^81^, GFP-labeled KobA (KobA-GFP), S-tag-labeled KobA (KobA-S), and S-tag-labeled KinA (KinA-S). Note that KobA can be pulled down with GFP-KinA-CBD but not with GFP-MyoE-GTD, while GFP-MyoE-GTD pulls down the MyoE-binding protein HmsV ^81^. KinA can also be pulled down with KobA-GFP or KobA-S. (B) A domain structure of KobA and a model of KinA-KobA interaction predicted by AlphaFold-3, followed by a predicted aligned error (PAE) plot. A lower PAE score, indicated by a darker blue color, indicates a better prediction with lower error. Note that both KinA and KobA form dimers. (C) Western analysis showing that KobA-HA (probed by the anti-HA antibody) is pulled down (indicated as “IP”) with KinA-GFP, GFP-KinA-CBD and GFP-KinA-CBD in a Δ*kinA* background. Uso1-GFP ^125^ serves as a negative control. The same blot also contains lanes with total protein extract indicated as “Input”. The GFP-fusion baits (probed by the anti-GFP antibody) are shown on another western blot on the right side. (D) Images of KobA-tdT in a wild-type *kinA* background and in the *kinA*-rigor (*kinA*^G97E^) background. Hyphal tip is indicated by a yellow arrowhead. Bar, 5 μm. Please see Figure S1 for kymographs and quantitative analyses of KobA-tdT in the wild-type *kinA* background. (E) Images of KobA-GFP in a wild-type *kinA* background and in the Δ*kinA* background. Hyphal tip is indicated by a yellow arrowhead. Bar, 5 μm. (F) Line scans of KobA-GFP fluorescence intensity in a wild-type *kinA* background and in the Δ*kinA* background. XY graphs with mean (solid lines) and SEM (shading) were generated by Prism 11. Between 0.130 μm and 5.200 μm from the hyphal tip, KobA-GFP intensity was significantly higher in wild type than that in the Δ*kinA* mutant (P < 0.0001) (two-way ANOVA with Šídák’s multiple comparisons test, *n* = 22 hyphae for wild type and *n* = 19 for Δ*kinA*). (G) A western blot showing that the protein level of KobA-GFP is significantly decreased in the Δ*kinA* mutant. (H) A quantitative analysis of KobA-GFP levels in the wild-type and Δ*kinA* strains. The intensity ratios of KobA-GFP to the loading control (indicated by Ponceau S staining) were calculated, and the ratios in the wild-type samples were set as 1. Scatter plots with mean and SD values as well as p values were generated by Prism 11 (Unpaired t test with Welch’s correction, two-tailed). (I) Colonies of a wild-type strain, a Δ*kinA* mutant, a Δ*kobA* mutant, and the Δ*kobA*, Δ*kinA* double mutant.

KobA is pulled down by either wild-type KinA or constitutively active KinA^K8^^95^* lacking the C-terminus involved in KinA autoinhibition ^71^ in Δ*hookA* cells missing the early endosome dynein adapter HookA ^72^ and in *alcA-nudK*^Arp1^ cells in which the expression of a key dynactin component Arp1 is repressed ^73^ (Table S1). These data suggest that the KobA-KinA association is independent of the KinA activation states or the cargo binding and activation status of dynein.

Our AlphaFold-3-based structural analysis predicts a direct interaction between a KinA dimer and a KobA dimer, with the interaction interface formed between the C-terminal part of KobA (aa167-414) and part of the KinA CBD (Figure 1B). That KinA and KobA interact via KinA CBD was also confirmed using GFP-tagged KinA CBD and HA-tagged KobA using western blots after pulldown experiments (Figure 1C). To observe KobA localization, we constructed a KobA-tdT fusion. KobA-tdT signals are enriched at the microtubule plus ends, as evidenced by the comet-like structures near the hyphal tip, resembling those formed by EB1, dynein or dynactin (Figure 1D) ^10,11,14,15,25,74^. In kymographs, the dynamic plus-end signals are obvious, and they move toward the hyphal tip with a speed similar to that of EB1-GFP (Figure S1). However, the signals that likely represent motor-driven movements (with a significantly higher speed) towards the plus ends are much fainter (Figure S1). To further demonstrate the KobA-KinA association in live cells, we used a *kinA*-rigor mutant (*kinA*^G97E^) where KinA should localize along microtubules ^27,30^. In the *kinA*^G97E^ background, KobA-tdT signals decorate microtubules (Figure 1D), consistent with KobA-KinA association. In addition, using a KobA-GFP construct, we found that the plus-end comet signals of KobA-GFP disappeared in the *kinA*-deletion mutant Δ*kinA* (Figure 1E, 1F) ^27^. Using western blot analyses, we found that the protein level of KobA-GFP is significantly decreased upon loss of KinA (Figure 1G, 1H), suggesting that KobA stability depends on KinA. Together, these results strongly support the notion that KobA is associated with KinA in live cells.

Homologs of KobA can be found in many species of Ascomycota, within the sub-phylum Pezizomycotina. Of note, a previously studied *Neurospora crassa* homolog of KobA, Ro16, has been shown to bind kinesin-1 CBD in yeast two-hybrid assays and participate in kinesin-1-mediated dynein localization (Ph.D. Thesis by T El Mellouki, 2016, University of Missouri–Kansas. https://mospace.umsystem.edu/server/api/core/bitstreams/43dca1ed-cdeb-4959-aa72-9e61672e2c49/content). Although sequence similarity is low between KobA and Ro16, the Spindly motif and the amino acid residues around it are highly conserved (Figure S2).

### KobA is required for KinA-mediated accumulation of dynein and dynactin at the microtubule plus end

To study KobA function in *A. nidulans*, we constructed a *kobA* deletion allele, Δ*kobA*. In colony growth tests, Δ*kobA* slightly impairs growth, although to a lesser extent than Δ*kinA* (Figure 1I). The Δ*kobA*, Δ*kinA* double mutant resembles the Δ*kinA* single mutant (Figure 1I), suggesting that KobA is in the same genetic pathway as KinA. To test if KobA plays a role in dynein localization, we introduced into the Δ*kobA* background a functional GFP-dynein fusion expressed from the endogenous dynein heavy chain *nudA* locus ^75^. In contrast to GFP-dynein in the wild-type background, which forms comet-like structures representing an accumulation at the microtubule plus ends near the hyphal tip ^10,11,75^, GFP-dynein in the Δ*kobA* background localizes along microtubule-like structures (Figure 2A, 2B), resembling its localization in the single Δ*kinA* mutant ^26^ and in the double Δ*kobA*, Δ*kinA* mutant backgrounds (Figure 2A, 2B). To rule out the possibility that these defects in dynein localization reflect that microtubules cannot extend normally towards the hyphal tip, we observed microtubules using GFP-labeled TubA (α tubulin) ^10,76^ and the mCherry-labeled plus-end-tracking protein EB1 ^25^ in the Δ*kobA* mutant. Overall, the microtubule network in the Δ*kobA* mutant looked similar to that in the wild type (Figure S3), strongly suggesting that the defect in dynein localization in the Δ*kobA* mutant is due to a defect in KinA-mediated transport of dynein rather than a gross abnormality of the microtubule network.

**Figure 2.**
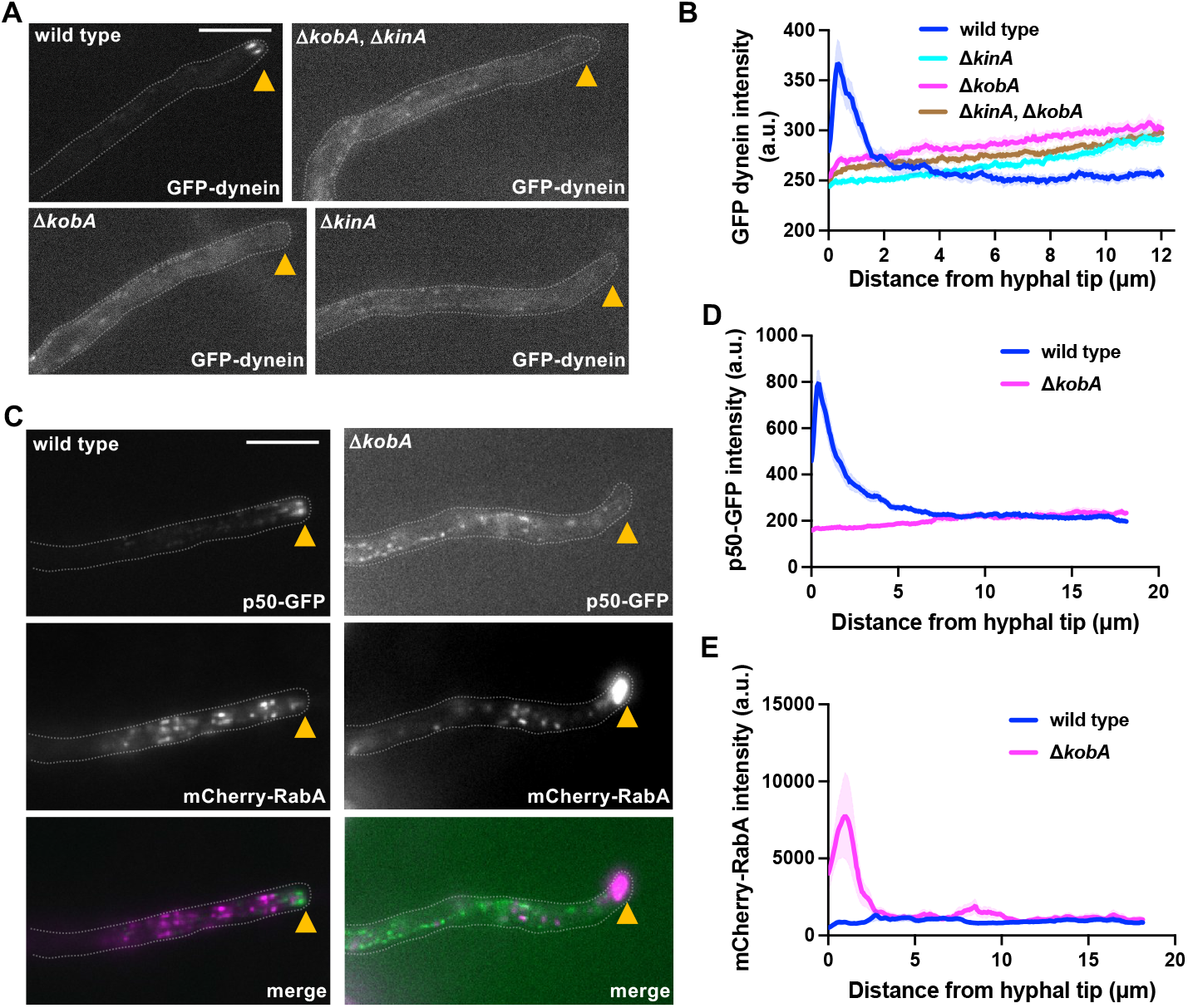
KobA is required for the accumulation of dynein and dynactin at the microtubule plus ends. (A) Microscopic images showing the distributions of GFP-dynein in wild type, Δ*kobA* mutant, Δ*kinA* mutant and Δ*kobA*, Δ*kinA* double mutant strains. Hyphal tip is indicated by a yellow arrowhead. Bar, 5 μm. (B) Line scans of GFP-dynein fluorescence intensity in wild type, Δ*kobA* mutant, Δ*kinA* mutant and Δ*kobA*, Δ*kinA* double mutant strains. XY graphs with mean (solid lines) and SEM (shading) were generated by Prism 11. Between 0.065 μm and 1.040 μm from hyphal tip, GFP-dynein intensity was significantly higher in wild type than that in the Δ*kobA* mutant, the Δ*kinA* mutant or the Δ*kobA*, Δ*kinA* double mutant (P < 0.0001) (two-way ANOVA with Tukey’s multiple comparisons test, *n* = 20 hyphae for all strains). (C) Microscopic images showing the distributions of dynactin p50-GFP as well as mCherry-RabA-labeled early endosomes (mCherry-RabA) in wild type and the Δ*kobA* mutant. Hyphal tip is indicated by a yellow arrowhead. (D) Line scans of p50-GFP fluorescence intensity in a wild-type background and in the Δ*kobA* background. XY graphs with mean (solid lines) and SEM (shading) were generated by Prism 11. Between 0 μm and 3.055 μm from hyphal tip, p50-GFP intensity was significantly higher in wild type than that in the Δ*kobA* mutant (P < 0.0001) (two-way ANOVA with Šídák’s multiple comparisons test, *n* = 12 hyphae for wild type and *n* = 10 for Δ*kobA*). (E) Line scans of mCherry-RabA (early endosome) fluorescence intensity in a wild-type background and in the Δ*kobA* background. XY graphs with mean (solid lines) and SEM (shading) were generated by Prism 11. Between 0 μm and 1.69 μm from hyphal tip, mCherry-RabA intensity was significantly higher in the Δ*kobA* mutant than that in wild type (P < 0.0001) (two-way ANOVA with Šídák’s multiple comparisons test, *n* = 12 hyphae for wild type and *n* = 10 for Δ*kobA*).

In addition to dynein, we examined the localization of dynactin and found that the Δ*kobA* mutant also exhibited a defect in the plus-end accumulation of the p50 subunit of the dynactin complex (Figure 2C, 2D). As the plus-end localization of dynactin is KinA-dependent ^14,28^, these data indicate that KobA cooperates with KinA also in the localization of dynactin. Kinesin-1-mediated dynein-dynactin localization to the hyphal tips facilitates the function of dynein in cargo transport; in *U. maydis* or *A. nidulans,* loss of kinesin-1 causes early endosomes, a dynein cargo, to abnormally accumulate at the hyphal tips where microtubule plus ends are located ^15,29,30^. Like in the Δ*kinA* mutant, early endosomes labeled with mCherry-RabA (or Rab5A ^77^) in the Δ*kobA* background abnormally accumulate at the hyphal tip (Figure 2C, 2E) ^26,29^.

### KobA specifically mediates dynein localization regardless of KinA or dynein activation states

Besides being required for dynein localization ^14^, kinesin-1s in filamentous fungi are required for transporting other cargoes toward the hyphal tip where microtubule plus ends are located ^27,78–80^. For example, KinA is required for transporting RabE-marked secretory vesicles to support hyphal tip growth ^27^. This transport is also mediated by myosin V (MyoV or MyoE) ^27,65,81^, such that the Δ*kinA*, Δ*myoV* double mutant is extremely sick and nearly non-viable ^27^. In contrast, the Δ*kobA*, Δ*myoV* double mutant formed a colony similar to that of Δ*myoV* (Figure 3A), strongly indicating that KobA is dispensable for the function of KinA in transporting secretory vesicles.

**Figure 3.**
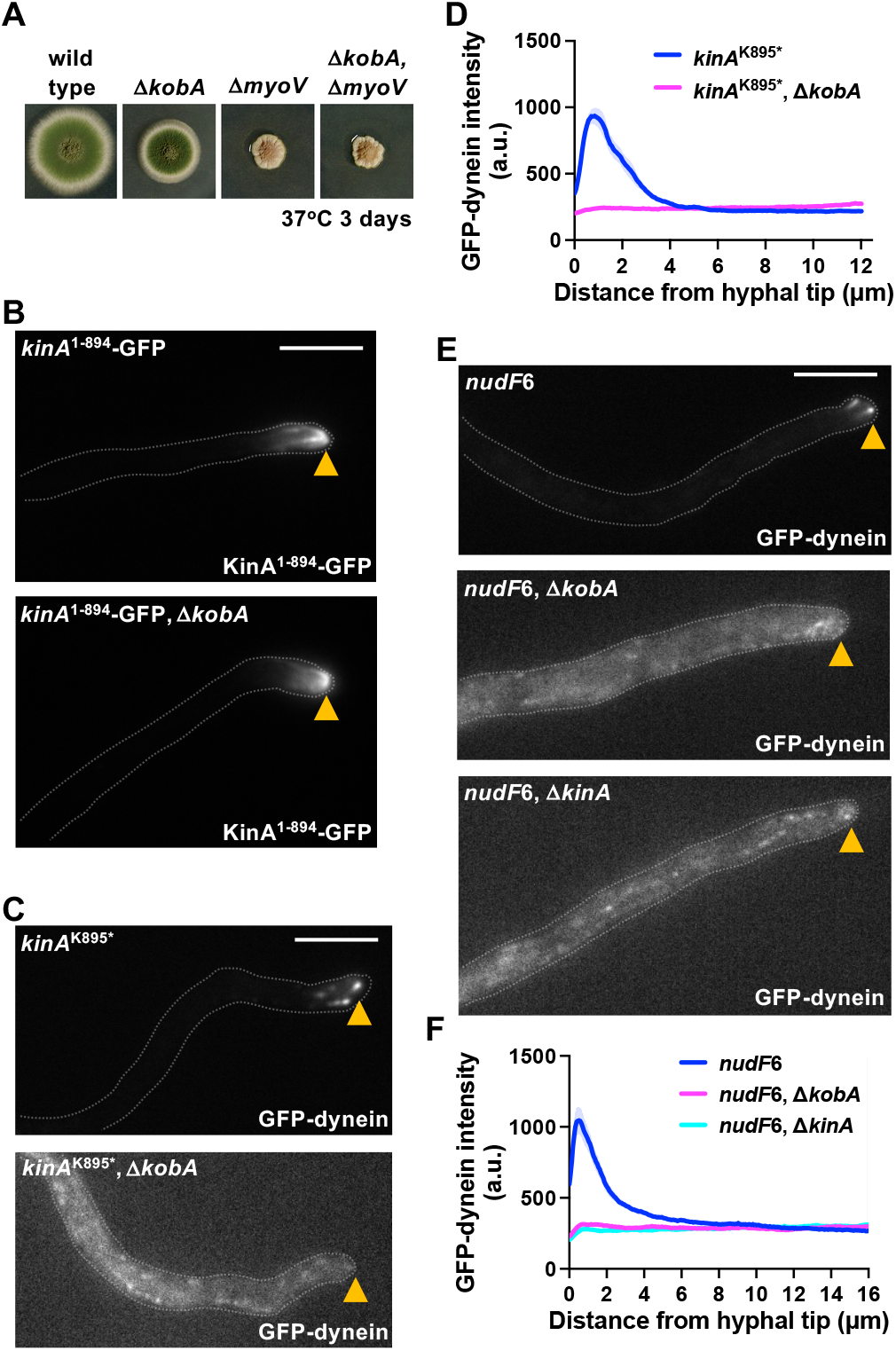
KobA is not required for the general motor activity of KinA, and its role in dynein localization is unrelated to the activation states of KinA or dynein. (A) Colony morphology of the Δ*myoV* mutant and the Δ*kobA*, Δ*myoV* double mutant. (B) Microscopic images showing the distributions of KinA^1-894^-GFP in wild type and the Δ*kobA* mutant background. Hyphal tip is indicated by a yellow arrowhead. Bar, 5 μm. (C) Microscopic images showing the distributions of GFP-dynein in the *kinA*^K895*^ single mutant and the Δ*kobA*, *kinA*^K895*^ double mutant. Hyphal tip is indicated by a yellow arrowhead. (D) Line scans of GFP-dynein fluorescence intensity in the *kinA*^K895*^ single mutant and the Δ*kobA*, *kinA*^K895*^ double mutant. XY graphs with mean (solid lines) and SEM (shading) were generated by Prism 11. Between 0.065 μm and 2.86 μm from hyphal tip, GFP-dynein intensity was significantly higher in *kinA*^K895*^ single mutant than that in the Δ*kobA*, *kinA*^K895*^ double mutant (P < 0.0001) (two-way ANOVA with Šídák’s multiple comparisons test, *n* = 23 hyphae for the *kinA*^K895*^ single mutant and *n* = 22 hyphae for the Δ*kobA*, *kinA*^K895*^ double mutant). (E) Microscopic images showing the distributions of GFP-dynein in the *nudF6* mutant, the *nudF*6, Δ*kobA* double mutant and the *nudF*6, Δ*kinA* double mutant. Hyphal tip is indicated by a yellow arrowhead. Bar, 5 μm. (F) Line scans of GFP-dynein fluorescence intensity in the *nudF*6 mutant, the *nudF*6, Δ*kobA* double mutant and the *nudF*6, Δ*kinA* double mutant. XY graphs with mean (solid lines) and SEM (shading) were generated by Prism 11. Between 0 and 3.835 μm from hyphal tip, GFP-dynein intensity was significantly higher in the *nudF*6 single mutant than in each of the double mutants (P < 0.0001), but there is no significant difference (p>0.05) between the *nudF*6, Δ*kinA* and *nudF*6, Δ*kobA* double mutants (two-way ANOVA with Tukey’s multiple comparisons test, *n* = 25 hyphae for *nudF*6, *n* = 23 hyphae for *nudF*6, Δ*kobA*, *n* = 22 hyphae for *nudF*6, Δ*kinA*).

Kinesin-1 is known to be regulated by autoinhibition ^37,39,42–44,51,52,54,82,83^, which involves a conserved C-terminal Ile-Ala-Lys (IAK) motif ^84,85^. While removing the IAK motif alone is insufficient to fully activate the motor in vitro ^39,54^, removing this motif or the *kinA*^K8^^95^* nonsense mutation truncating the protein within this motif disrupts KinA autoinhibition in *A. nidulans* ^71^. In contrast to KinA-GFP that shows diffuse signals with a slight hyphal-tip enrichment, KinA^1-8^^94^-GFP shows a strong accumulation at the hyphal tip with some signals clearly decorating microtubules near plus ends (Figure 3B) ^71^. In the Δ*kobA* background, KinA^1-8^^94^-GFP still shows a strong accumulation at the hyphal tip, suggesting that the general motor activity of KinA is not abolished in the Δ*kobA* mutant (Figure 3B). However, although GFP-dynein can still accumulate at the hyphal tip in the *kinA*^K8^^95^* background ^71^, dynein in the Δ*kobA*, *kinA*^K8^^95^* double mutant background localizes along microtubule-like structures (Figure 3C, 3D). Thus, KobA is still required for KinA-mediated dynein accumulation at the microtubule plus ends even when KinA autoinhibition is disrupted.

One possibility was that KobA keeps dynein in an inactive conformation during its plus-end-directed transport. In this scenario, the absence of KobA would result in premature activation of dynein before it arrives at the plus ends of microtubules, consequently resulting in a tug-of-war between dynein and KinA, which would prevent dynein’s plus-end accumulation. In *A. nidulans, nudF* encodes the LIS1 (Lissencephaly-1 protein) homolog NudF ^86^, and dynein in the *nudF*6 (lis1) (or *nudF*^L304S^) mutant is inactive or autoinhibited ^87^, consistent with other recent studies ^57,88–100^. In the *nudF6* temperature-sensitive mutant grown at the restrictive temperature of 37°C, dynein fails to be activated by the dynein adapter HookA due to its autoinhibited state ^87^, thereby accumulating at the microtubule plus ends and resulting in strong GFP-dynein signals at the hyphal tip (Figure 3E, 3F). However, in cells of the *nudF*6, Δ*kobA* double mutant and the *nudF*6, Δ*kinA* double mutant, GFP-dynein shows a localization pattern consistent with microtubule decoration (Figure 3E, 3F). This result strongly suggests that even when dynein is inactive, KobA is still needed for KinA’s function in transporting dynein to the plus end.

### KobA facilitates the KinA-dynactin-dynein association

Previously, we have shown that the dynactin complex enhances the physical interaction between KinA and dynein ^64^, suggesting that dynein and dynactin are bound to KinA to be transported to the plus end together. Consistent with this idea, the microtubule plus-end accumulation of dynactin in the Δ*kobA* mutant is also significantly reduced (Figure 2C, 2D). KobA contains a Spindly motif that is known to bind the pointed-end protein p25 of the dynactin complex ^66–69^. In an AlphaFold-3-based structural prediction on KobA-dynactin interaction, the KinA-KobA complex is predicted to interact with part of the dynactin complex including the Arp1 mini-filament and the pointed-end sub-complex, with the Spindly motifs of the KobA dimer being close to p25 (Figure 4A, 4B).

**Figure 4.**
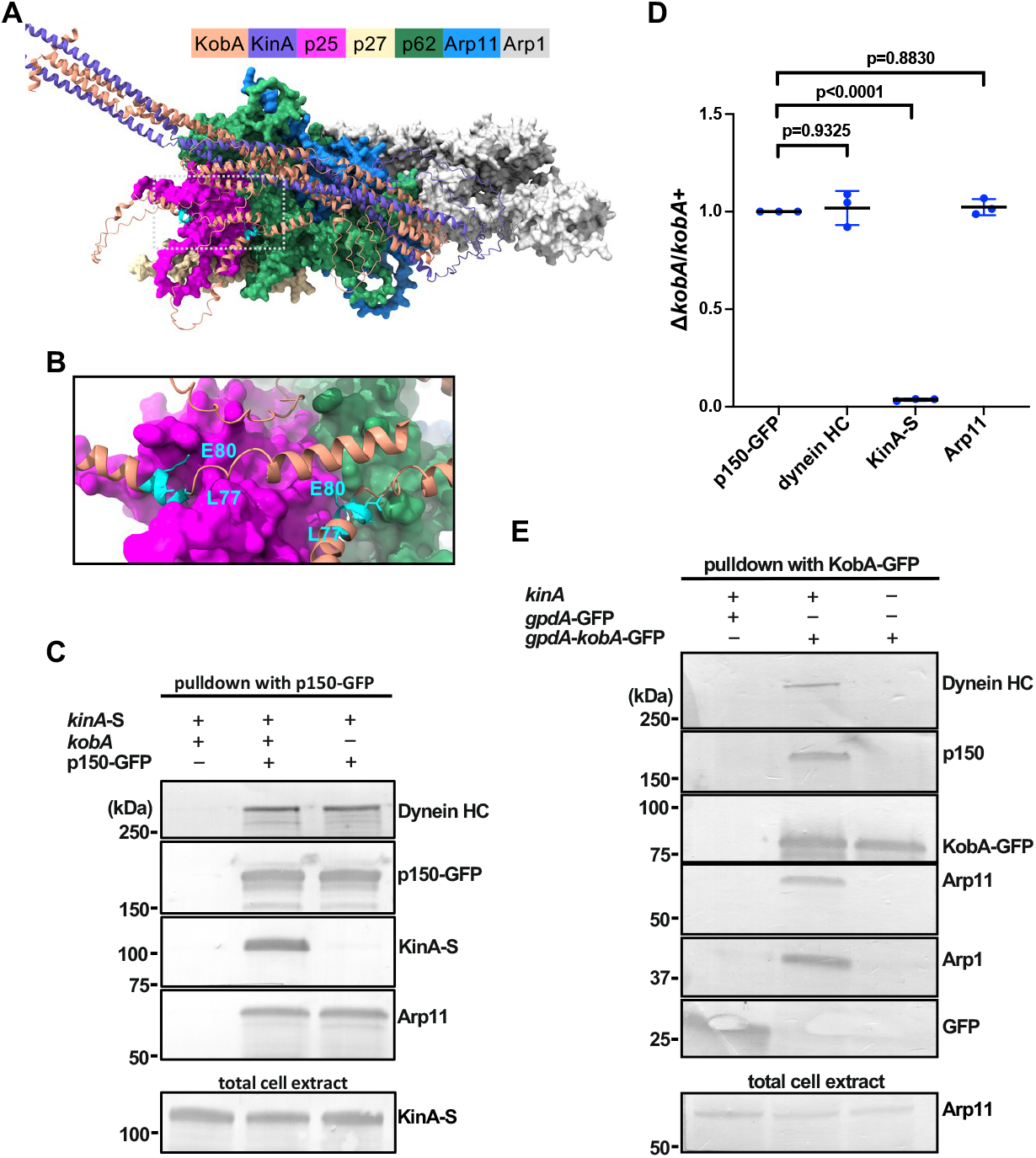
KobA is required for the interaction between KinA and dynactin-dynein. (A) A model of the KinA-KobA-dynactin interactions predicted by AlphaFold-3. (B) A zoom-in of the p25-KobA interactions. The two “Spindly motifs” of the KobA dimer are colored in cyan (L77 and E80 are highlighted), although KobA is colored in light brown in the model. (C) Western blots showing that the amount of KinA-S pulled down with p150-GFP is dramatically reduced in the Δ*kobA* (or *kobA* -) background. A strain containing KinA-S alone without p150-GFP (or p150-GFP -) is used as a negative control (the leftmost lane). (D) A quantitative analysis on the effect of *ΔkobA* on the amounts of dynein HC, KinA-S and Arp11 pulled down with p150-GFP. The ratios of the intensity values of *ΔkobA* to those of *kobA+ (*wild-type *kobA)* are shown for each component, and these ratios are relative to the ratio for p150-GFP that was set as 1. Scatter plots with mean and SD values as well as p values were generated by Prism 11 (Ordinary one-way ANOVA with Dunnett’s multiple comparison test). (E) Western blots showing that dynein and dynactin are pulled down with overexpressed KobA-GFP (*gpdA*-KobA-GFP) in the wild-type background but not in the Δ*kinA* (or *kinA* -) background. A strain containing overexpressed GFP (*gpdA*-GFP) is used as a negative control (the leftmost lane).

We tested experimentally if KobA plays a role in the KinA-dynactin association. Specifically, we performed pulldown experiments using strains containing p150-GFP and KinA-S ^64^, with or without KobA (Δ*kobA*). A strain containing KinA-S alone was used as a negative control. We found that while p150-GFP pulled down KinA-S in the strain with wild-type *kobA*, the amount of KinA-S pulled down with p150-GFP was diminished markedly in the Δ*kobA* mutant background (Figure 4C, 4D). This result suggests that KobA helps link KinA to dynactin-dynein.

We also tested whether the KobA-dynactin interaction depends on KinA. Because the protein level of KobA-GFP is significantly decreased in the Δ*kinA* mutant (Figure 1G, 1H), we generated a strain containing a *gpdA*-KobA-GFP allele, which drives overexpression of KobA-GFP under the control of the *gpdA* (*g*lyceraldehyde-3-*p*hosphate *d*ehydrogenase) promoter ^101,102^. In a wild-type *kinA* background, overexpressed KobA-GFP pulled down both dynein and dynactin (Figure 4E). However, in the Δ*kinA* background, we could hardly detect any dynein-dynactin pulled down with KobA-GFP (Figure 4E). Thus, KobA and KinA mutually depend on each other to link them to dynactin-dynein.

### The Spindly motif of KobA plays an important role in the KinA-dynactin interaction

We next sought to directly test the role of the Spindly motif of KobA in the KinA-dynactin interaction. We edited *kobA* to introduce two missense mutations resulting in a Leu77Ala, Glu80Ala (*kobA*^L77A,E80A^) double substitution in the motif. To facilitate the genetic, imaging and biochemical analyses of the mutant, we linked mCherry at the C-terminus of KobA^L77A,E80A^ to generate the *kobA*^L77A,E80A^-mCherry allele. As a control, we also generated a *kobA*-mCherry allele. We introduced the *kobA*^L77A,E80A^-mCherry and the *kobA*-mCherry alleles separately into the background of p150-GFP, KinA-S for pulldown analyses. In these experiments, we also included the Δ*kobA* mutant and the Δp25 mutant for comparisons. We found that the *kobA*^L77A,E80A^-mCherry allele significantly reduced the amount of KinA-S pulled down by p150-GFP, although the extent of the reduction is smaller compared to that caused by Δ*kobA* (Figure 5A, 5B). The Δp25 mutation also significantly reduced the amount of KinA-S pulled down with p150-GFP, which is consistent with our previous results ^64^. These data provide strong support for the AlphaFold-3-based structural prediction that the KinA-KobA-dynactin association involves the direct interaction between the KobA Spindly motif and dynactin pointed end, although other regions of KobA may further enhance the association.

**Figure 5.**
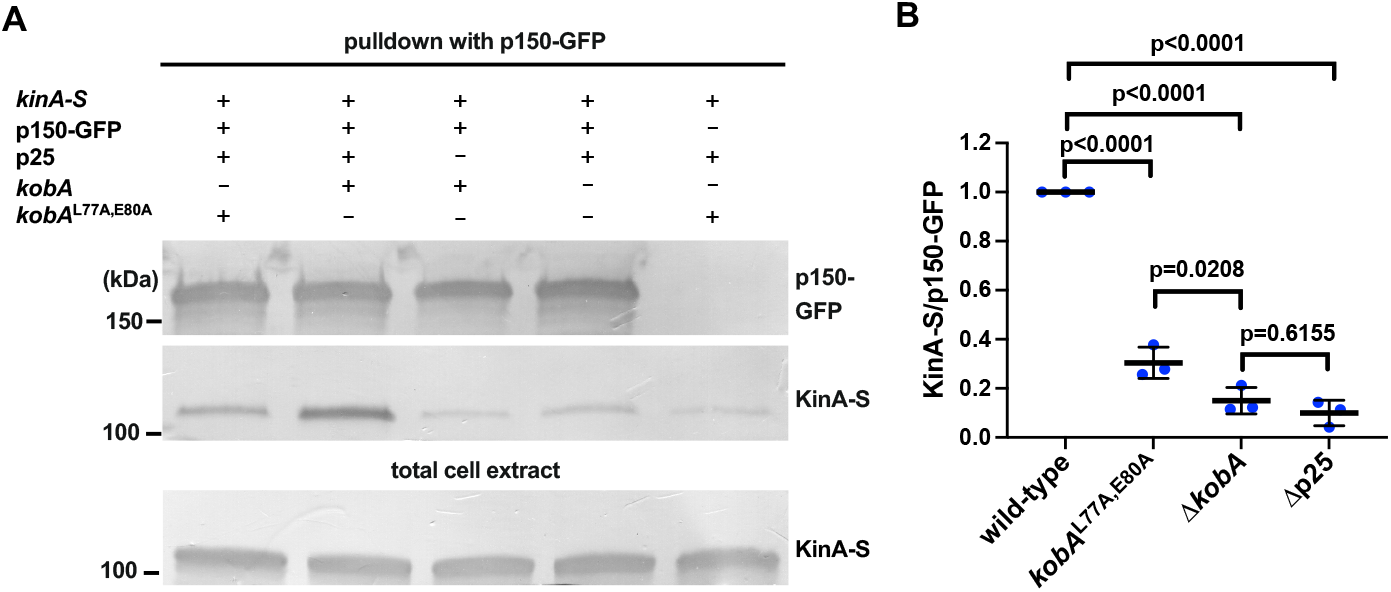
The Spindly motif of KobA enhances the KinA-dynactin interaction. (A) Western blots showing that the amount of KinA-S pulled down with p150-GFP is obviously reduced in the *kobA*^L77A,E80A^-mCherry, Δ*kobA* and Δp25 backgrounds compared to that in the *kobA*-mCherry background without the *kobA*^L77A,E80A^ Spindly-motif mutations. The strain containing *kobA*^L77A,E80A^-mCherry and KinA-S but not p150-GFP was used as a negative control (the rightmost lane). (B) A quantitative analysis on the effects of *kobA*^L77A,E80A^-mCherry, Δ*kobA* and Δp25 on the amounts of KinA-S pulled down with p150-GFP. Note that the background intensity values of KinA-S in the negative control were subtracted from the KinA-S signal intensity in other lanes. The ratios of the intensity values of KinA-S to p150-GFP are shown and these ratios are relative to the ratio for the *kobA*-mCherry-containing strain (called wild-type in this figure) that was set as 1. Scatter plots with mean and SD values as well as p values were generated by Prism 11 (Ordinary one-way ANOVA with Dunnett’s multiple comparison test).

Consistent with a role of the Spindly motif of KobA in enhancing the KinA-dynactin interaction, we found that *kobA*^L77A,E80A^-mCherry significantly decreased the plus-end accumulation of GFP-dynein, and led to a localization pattern of GFP-dynein consistent with microtubule decoration (Figure 6A, 6B). To examine whether the Spindly motif mutations affect the KobA-KinA interaction in vivo, we examined *kobA*^L77A,E80A^-mCherry and *kobA*-mCherry in the *kinA*^1-894^–GFP background where KinA autoinhibition is disrupted ^71^. In the Spindly-motif mutant, KobA is still able to co-localize with KinA at the hyphal tip (Figure S4), although the hyphal-tip intensity of KinA-GFP or KobA-mCherry is lowered, possibly because cargo interaction (in the case the cargo is dynein-dynactin) could further activate KinA that lacks the C-terminal sequence for autoinhibition (Figure S4) ^39,54^. The ratio of KobA-mCherry to KinA-GFP signal intensity along the hypha is not affected by the *kobA*^L77A,E80A^ mutations, suggesting that the Spindly motif of KobA is unlikely to affect the KobA-KinA interaction, but rather, it is involved in the KobA-dynactin interaction as predicted by AlphaFold-3.

**Figure 6.**
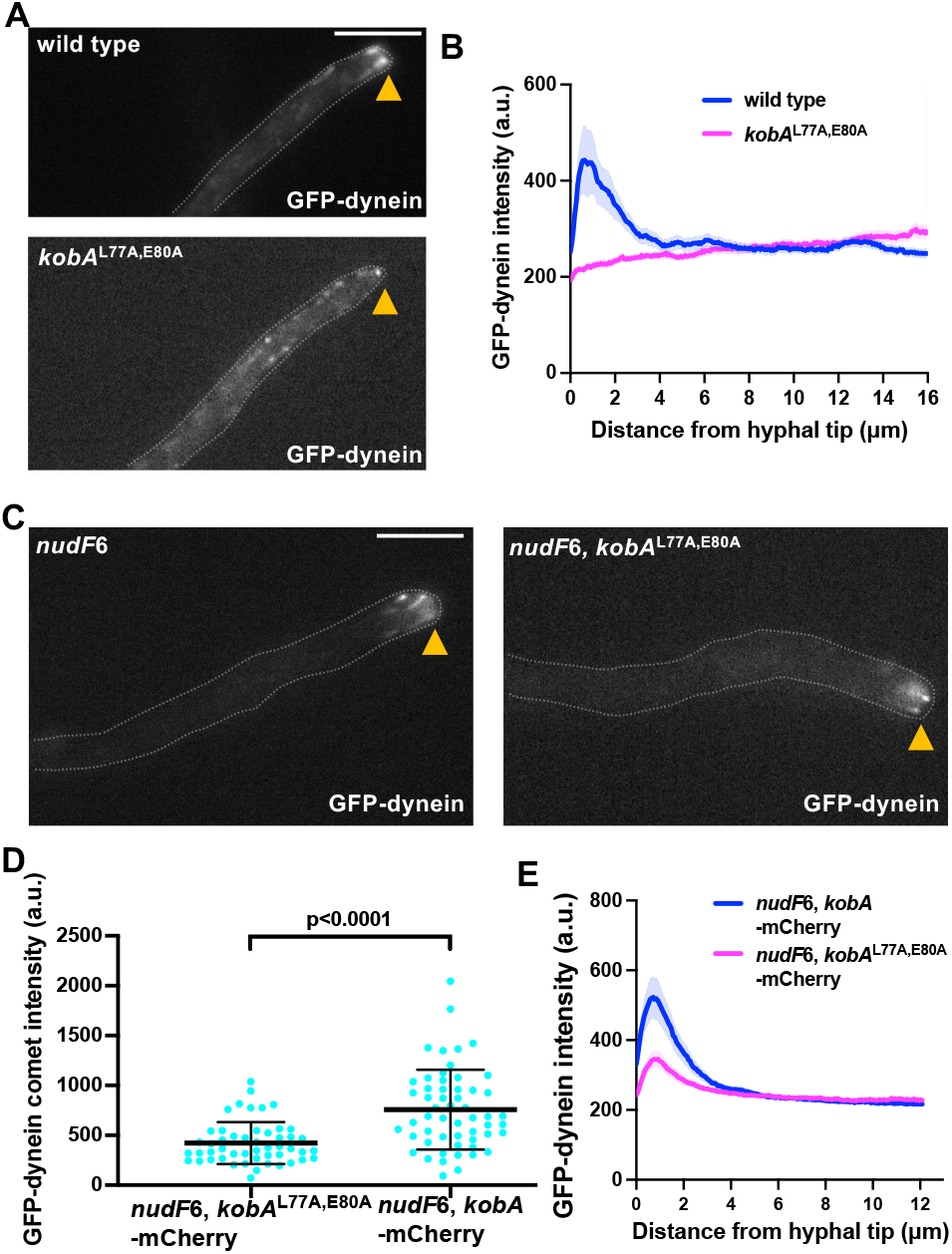
The Spindly motif is required for the plus-end localization of dynein. (A) Microscopic images showing the distributions of GFP-dynein in wild type and the *kobA*^L77A,E80A^ mutant. Hyphal tip is indicated by a yellow arrowhead. Bar, 5 μm. (B) Line scans of GFP-dynein fluorescence intensity in wild type and the *kobA*^L77A,E80A^ mutant. XY graphs with mean (solid lines) and SEM (shading) were generated by Prism 11. Between 0.26 and 1.885 μm from hyphal tip, GFP-dynein intensity in wild type was significantly higher than that in the *kobA*^L77A,E80A^ mutant (p<0.0001, two-way ANOVA with Šídák’s multiple comparisons test, *n* = 22 hyphae for wild type, *n* = 21 hyphae for *kobA*^L77A,E80A^). (C) Microscopic images showing the distributions of GFP-dynein in the *nudF*6, *kobA*-mCherry and the *nudF*6, *kobA*^L77A,E80A^-mCherry strains. Hyphal tip is indicated by a yellow arrowhead. (D) A quantitative analysis of plus-end dynein accumulation (or GFP-dynein comet intensity) in the *nudF*6, *kobA*-mCherry and the *nudF*6, *kobA*^L77A,E80A^-mCherry strains. Scatter plots with mean and SD values were generated by Prism 11, and the p value was generated by Mann-Whitney test (unpaired, two-tailed). (E) Line scans of GFP-dynein fluorescence intensity in the *nudF*6, *kobA*-mCherry and the *nudF*6, *kobA*^L77A,E80A^-mCherry strains. XY graphs with mean (solid lines) and SEM (shading) were generated by Prism 11. Between 0.130 and 1.495 μm from the hyphal tip, GFP-dynein intensity in *nudF*6, *kobA*-mCherry was significantly higher than that in the *nudF*6, *kobA*^L77A,E80A^-mCherry mutant (p<0.0001, two-way ANOVA with Šídák’s multiple comparisons test, *n* = 30 hyphae for *nudF*6, *kobA*-mCherry, *n* = 29 hyphae for *nudF*6, *kobA*^L77A,E80A^-mCherry).

Finally, we examined the effect of the Spindly motif mutations *kobA*^L77A,E80A^-mCherry in the *nudF*6 (lis1) (*nudF*^L304S^) mutant background where dynein is inactive ^87^. In the *nudF*6, *kobA*^L77A,E80A^-mCherry double mutant, GFP-dynein can still be observed to accumulate at the plus end, but the intensity of this accumulation is significantly decreased compared to that of the *nudF*6, *kobA-*mCherry strain (Figure 6C, 6D, 6E). Thus, the Spindly motif of KobA plays an important role in the plus-end accumulation of dynein even when dynein is inactive, although this role becomes less obvious in this case possibly because the plus-end dynein accumulation is significantly enhanced by the *nudF*6 mutation (Figure S5).

## Discussion

In elongated cells such as neurons and fungal hyphae, cytoplasmic dynein and one of its activators, the dynactin complex, must first be carried outward to microtubule plus ends by kinesin-1 before dynein-dynactin can be activated by cargo adapters to drive retrograde transport ^14,15,19,21,35,36^. How kinesin-1 recognizes and transports dynein and dynactin remains a major mechanistic puzzle across cell types. Here we discovered a kinesin-1-binding protein in *A. nidulans*, KobA, which plays a critical role in the KinA kinesin-1-mediated plus-end accumulation of dynein and dynactin. KobA is a coiled-coil protein that binds to the C-terminal cargo-binding domain of kinesin-1 (KinA). Crucially, KobA is dispensable for KinA’s conventional cargo transport (such as secretory vesicle delivery alongside myosin-V ^27^), indicating that KobA functions specifically as a dynein-dynactin adapter rather than a general kinesin cofactor.

Strikingly, KobA harbors a conserved Spindly motif within its disordered N-terminus. Spindly motifs are widely known as signatures of dynein-activating adapters that engage the dynactin pointed-end protein p25 ^66–70^. Note that the Spindly motifs found in various dynein activating adapters have a consensus sequence of JxxEΦ where J is leucine or isoleucine, x can be any amino acid and Φ can be any small hydrophobic amino acid ^69^. The Spindly motif in KobA, LADEL, is particularly similar to the Spindly motif LAEEL of the dynein activating adapter HAP1 (Huntingtin-Associated Protein 1), with two negatively charged residues ^103^. HAP1 interacts with Huntingtin, whose Huntington’s disease-causing polyglutamine (polyQ) expansion increases HAP1 and kinesin-1 recruitment to BDNF endosomes ^104^. HAP1 binds to kinesin-1 ^2,105^, and specifically, HAP1 is a KIF5 kinesin-1 adapter required for KIF5-mediated transport of GABA(A) receptors to the synapse ^105^. HAP1’s Spindly motif is important for dynein activation ^103^, but it is not known to have any role in kinesin-1-mediated transport of dynein in neurons. We show that KobA in *A. nidulans* repurposes the same class of motif to recruit dynactin-dynein as kinesin-1 cargo.

Supported by AlphaFold-3 modeling, our biochemical pulldown results indicate that KobA and KinA depend mutually on each other to form a complex with dynactin and dynein. Point mutations disrupting the KobA Spindly motif (L77A, E80A) significantly weaken the KinA–dynactin interaction, resulting in a clear defect in the localization of dynein. Specifically, in the *kob*^AL77A,E80A^ mutant, dynein is localized along microtubules rather than accumulating at the plus ends. Consistent with our current structural prediction and the prior structural work demonstrating the interaction between the Spindly motif and p25 of dynactin ^57,66,69,70^, we have also found that the KinA-dynactin interaction is significantly disrupted in the Δp25 mutant. This result is consistent with our previous pulldown data suggesting a role of p25 in the KinA-dynactin interaction as well as the live-cell imaging data showing that p25 plays a critical role in the plus-end accumulation of dynactin ^64^. Specifically, in the Δp25 mutant, dynactin is localized along microtubules instead of accumulating at the plus ends ^64^. Given that KobA and the dynein-activating adapter both use the Spindly motif to interact with dynactin p25, it is possible that KobA binding to p25 may prevent premature dynein activation caused by binding of the dynein-activating adaptors. Otherwise, activated dynein-dynactin may overpower kinesin-1 to inhibit its plus-end motility ^106^. In this context, we find that inactive dynein in the *nudF*6 background can still accumulate at the plus ends although the accumulation is significantly decreased in the *kob*^AL77A,E80A^ mutant background. Because the Spindly motif of KobA has a bridging function, it is hard to dissect its additional role in preventing premature dynein activation, but such a role is not excluded and would deserve to be investigated in the future. In addition, the coiled-coil region of KobA is likely to be also involved in linking KinA to dynactin-dynein as predicted (Figure 4A), and the detailed interactions will need to be further dissected.

KobA sequence homologs are found in many species of Ascomycota, within its largest sub-phylum Pezizomycotina that has ∼85,000 species ^107^, including dangerous human or plant pathogens such as *Aspergillus fumigatus*, *Magnaporthe oryzae* and *Fusarium graminearum.* In both *A. nidulans* and *A. fumigatus*, defects in microtubule-based transport are also linked to alterations in medically relevant secondary metabolites ^108^. Sequence conservation among KobA homologs in Pezizomycotina, including *N. crassa* (Ro16) https://mospace.umsystem.edu/server/api/core/bitstreams/43dca1ed-cdeb-4959-aa72-9e61672e2c49/content), *Magnaporthe oryzae* (synonym of *Pyricularia oryzae*), and *Fusarium graminearum*, is not high, but all these homologs have a Spindly motif LADEL (Figure S2). In the filamentous fungus *Ustilago maydis*, a plant pathogen not within Ascomycota but within Basidiomycota, kinesin-1 is also required for dynein accumulation at microtubule plus ends ^15^. Using a structure-based search, we found a *U. maydis* protein (UM04544) that may interact with *U. maydis* kinesin-1 (Kin-1) as predicted by AlphaFold-3 (Figure S6). UM04544 does not show strong sequence homology with KobA, but it contains a putative Spindly motif LEDEL. Thus, while obvious sequence homologs of KobA can only be found in Pezizomycotina, we cannot exclude the possibility that there are functional homologs of KobA in higher eukaryotes.

That the KobA-dynactin interaction is required for the plus-end accumulation of dynein suggests that dynein and dynactin are transported to the plus ends together in *A. nidulans*. In neurons, dynein and dynactin are both transported anterogradely, but evidence suggests that they may be transported separately ^18,20^. While it is unclear how dynactin is transported to the plus end by kinesin-1, the kinesin-1-mediated dynein transport in neurons may involve the direct interaction between dynein intermediate chains and kinesin light chains ^19,60^. Kinesin-1 light chains contain TPR domains and their functions are implicated in both reinforcing kinesin-1 autoinhibition and helping kinesin-1 bind cargoes ^37,39,52,54,109–115^. KobA does not contain TPR domains and is structurally distinct from kinesin light chains. We do not have any evidence to suggest that KobA would reinforce KinA autoinhibition prior to binding to the cargo, which is the dynein-dynactin complex. Instead, KobA is required for KinA-mediated dynein transport even when KinA is constitutively active. Because fungi lack kinesin light chains ^61,62^, the discovery of KobA demonstrates a novel evolutionary strategy where a dedicated coiled-coil adapter utilizes dynactin pointed-end recognition motifs to achieve plus-end dynein delivery.

Among the dynein cargo adapters, JIP3 was the only one that has been implicated in kinesin-1-based transport of dynein to the microtubule plus ends in *C. elegans* ^31^. JIP3 is known as a dynein activator ^56–59^, and it binds to the CC4 domain of the mammalian kinesin-1 KIF5B ^116^, which is similar to the region of KinA that KobA binds to. JIP3 uses different sites to bind kinesin-1 and dynein, which may allow JIP3 to bind kinesin-1 and dynein-dynactin at the same time ^116^. However, the Spindly motif of JIP3 has never been implicated in the plus-end-directed transport of dynactin-dynein. Other dynein cargo adapters may also bind kinesins (including and beyond kinesin-1s) and the directionality of the complex is regulated in different ways ^40,51,55,117–124^. Although KobA is a coiled-coil protein with a Spindly motif, we do not think KobA is a dynein-activating adapter. From our pulldown assays, it is obvious that the KobA-KinA interaction is much stronger and easier to detect than the KobA-dynactin-dynein interaction, and the KinA-KobA interaction is independent of dynactin (Table S1). It is possible that the KinA-KobA complex interacts dynamically with the inactive dynein-dynactin complex along microtubules and brings them to the plus ends where dynein-dynactin can interact with the early endosome cargo bound to the Hook adapters, followed by adapter-mediated dynein activation to move early endosomes toward the minus ends of microtubules (Figure 7).

**Figure 7.**
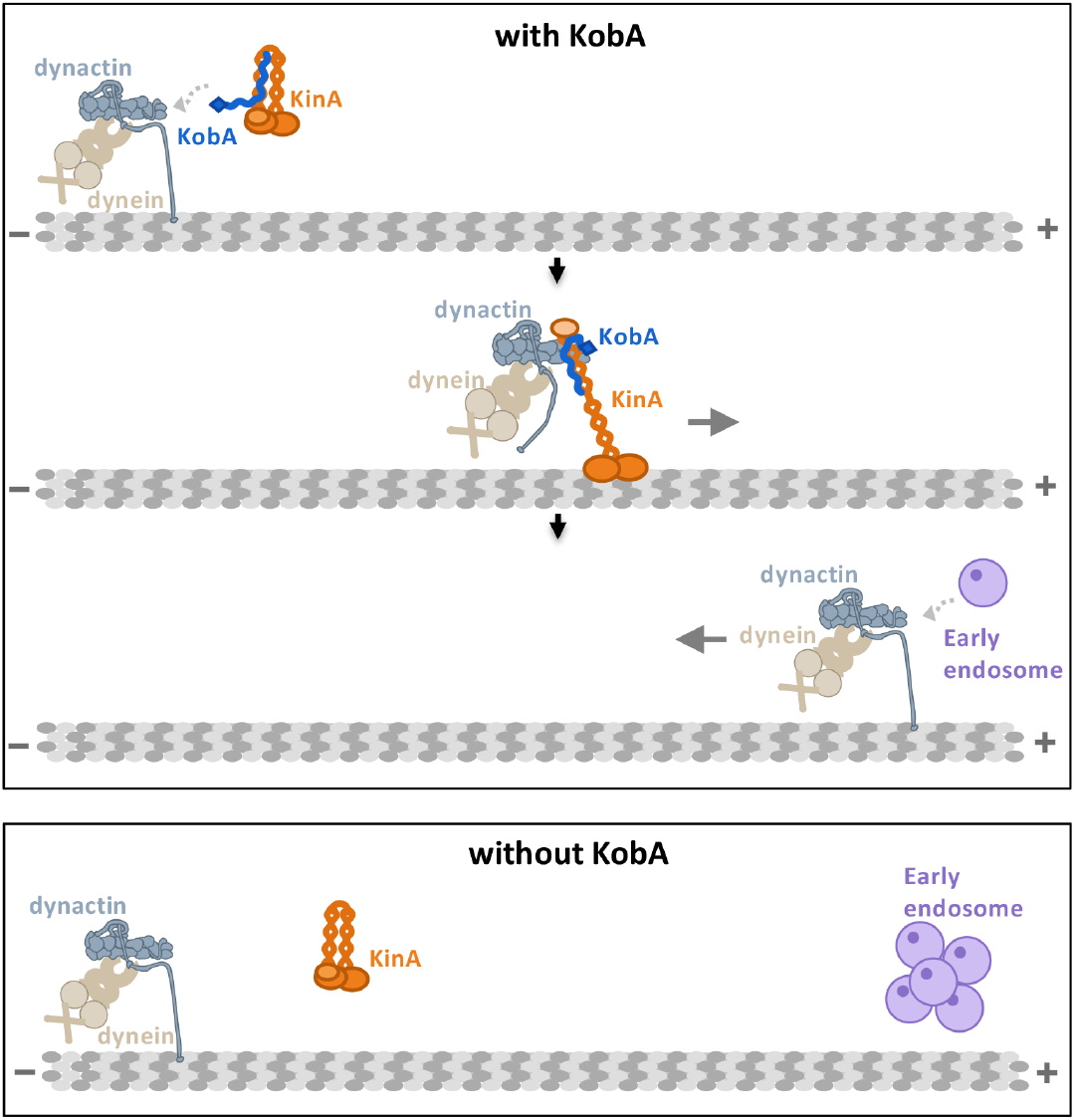
A model of how KobA bridges KinA with dynactin-dynein and moves them to the plus ends of microtubules. Before dynein-dynactin interacts with the kinesin-1 KinA, KinA is in an autoinhibited conformation, but KobA is able to bind KinA, as evidenced by the pulldown data indicating that the KobA-KinA interaction is independent of either the KinA-autoinhibition states or Arp1 of dynactin (Table S1). Dynein and dynactin can interact normally with each other in the absence of KobA-KinA as shown by the pulldown data (Figure 4C, 4D). The microtubule-binding domain of p150 is important for dynein-dynactin to interact with microtubules in the absence of KinA. This notion is based on our previous data that while wild-type p150 localizes along microtubules in the absence of KinA, p150 missing the microtubule-binding domain is diffuse in the cytoplasm ^28^. When a KobA-KinA complex interacts with dynein-dynactin, the KinA-KobA-dynactin-dynein complex moves toward the microtubule plus end. At the plus end, dynactin-dynein engages the early endosome cargoes bound to the Hook adapters, which activate dynein-dynactin and cause them to move toward the minus ends. Without KobA, dynactin-dynein fails to be transported to the plus ends, and early endosomes accumulate abnormally near the plus ends.

## Acknowledgements

We thank Berl Oakley, Reinhard Fischer, John Salogiannis, Samara Reck-Peterson, Martin Egan, Herb Arst and Steve Osmani for Aspergillus strains. This work was funded by the National Institutes of Health R35GM140792 (to Xin Xiang); a Department of War #OSD(HA).2022ICD.WBH-1 Joint DOTML-PF Warfighter Brain Health grant to USUHS Department of Biochemistry and Molecular Biology; grants TED2021-129607B-I00 funded by MCIN/AEI/10.13039/501100011033 and by the “European Union Next Generation EU/PRTR”; by grant PID2021-124278OB-I00 funded by MCIN/AEI/10.13039/501100011033/FEDER, UE and “ERDF A way of making Europe”; and by grant PID2024-1-155595NB-I00 funded by MCIN/AEI/10.13039/501100011033/FEDER, UE to Eduardo A. Espeso and Miguel A. Peñalva. Disclaimer: The opinions and assertions expressed herein are those of the author(s) and do not necessarily reflect the official policy or position of the Uniformed Services University or the Department of War.

---

**Figure S1.**
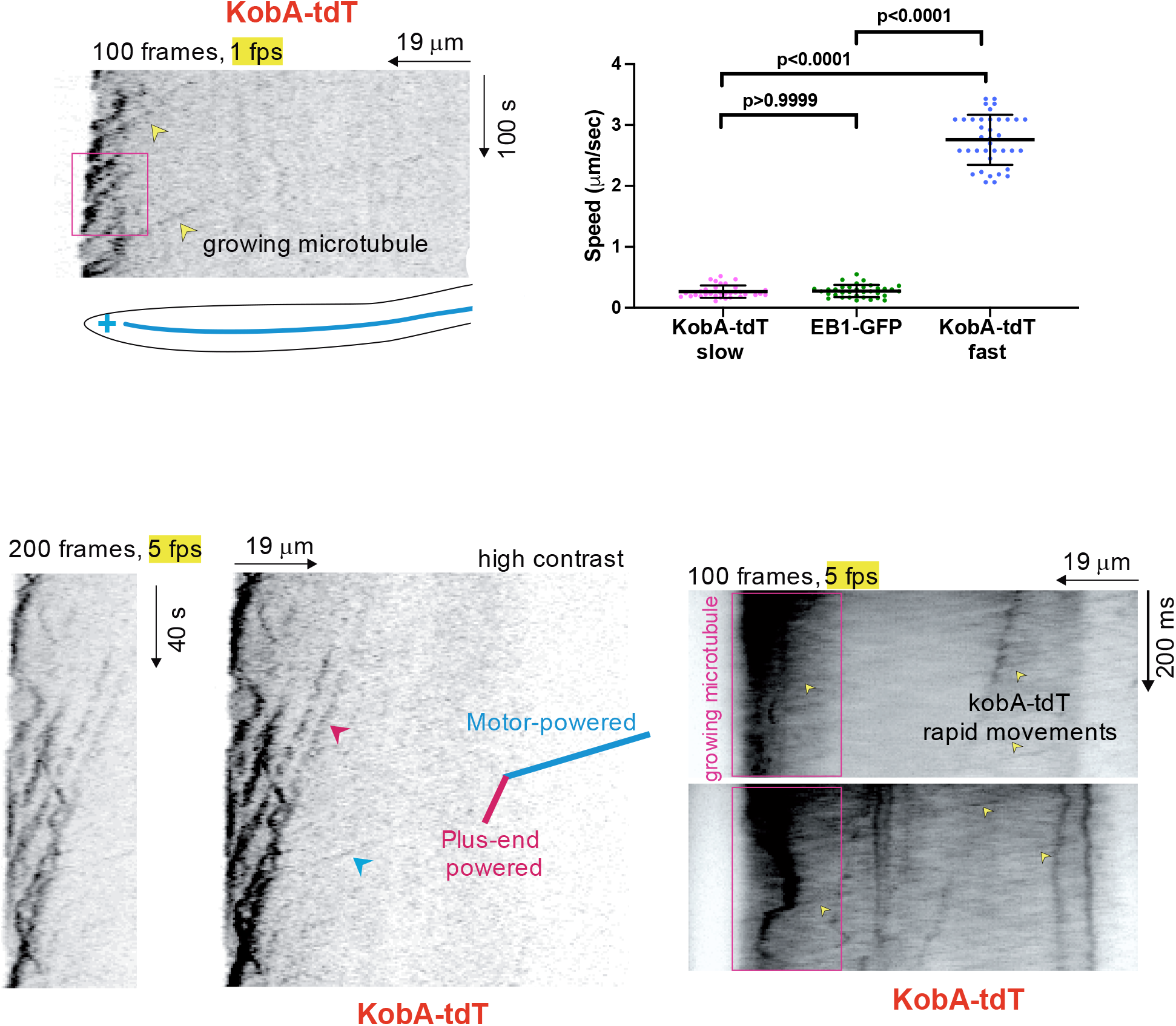
Kymographs and quantitative analysis of the KobA-tdT signals in a wild-type background (linked to Figure 1D). In all these kymographs, the hyphal tips with the growing microtubule ends (the plus ends) are on the left side. Two kinds of movements are revealed: the slow movements represent the dynamic microtubule plus-end-associated signals near the hyphal tip; and the rapid movements represent those driven by a plus-end-directed motor, most likely KinA. The slow movements have speeds similar to those of the end-binding protein EB1-GFP, suggesting that the speeds represent those of microtubule growth. Signals representing the motor-driven movements are much fainter than those associated with the growing plus ends and move with significantly higher speeds than those of EB1-GFP. For the quantitative analysis, scatter plots with mean and SD values as well as p values were generated by Prism 11 (Kruskal-Wallis ANOVA test with Dunn’s multiple comparisons test, unpaired).

**Figure S2.**
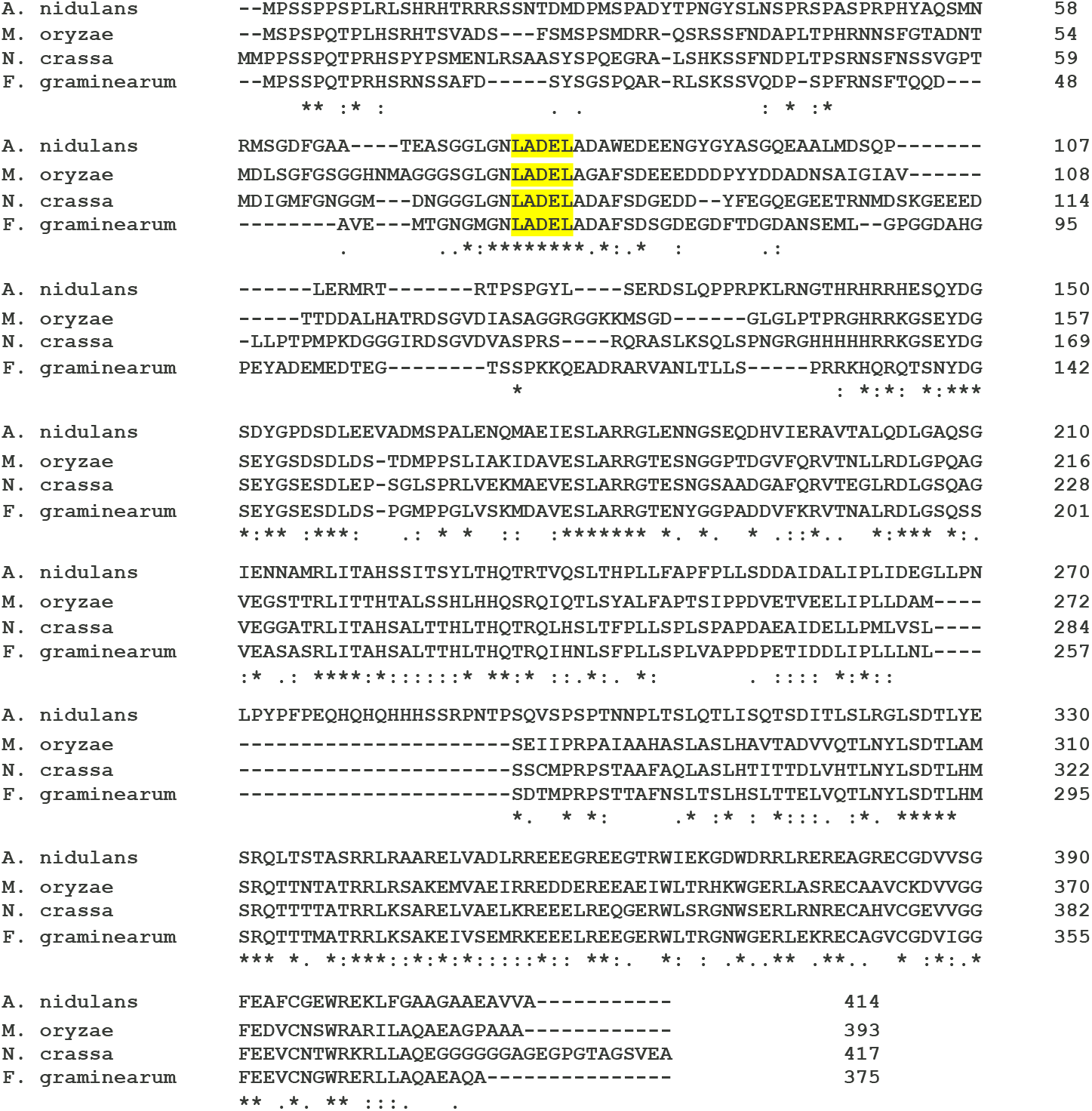
Multiple sequence alignment of KobA (An2744) from *Aspergillus nidulans* and its orthologs from *Magnaporthe oryzae* (synonym of *Pyricularia oryzae*), *Neurospora crassa*, and *Fusarium graminearum*. The alignment was done using Clustal Omega (1.2.4) at website https://www.ebi.ac.uk/jdispatcher/msa/clustalo. Residues that are identical (*), strongly similar (:) or weakly similar (.) are indicated.

**Figure S3.**
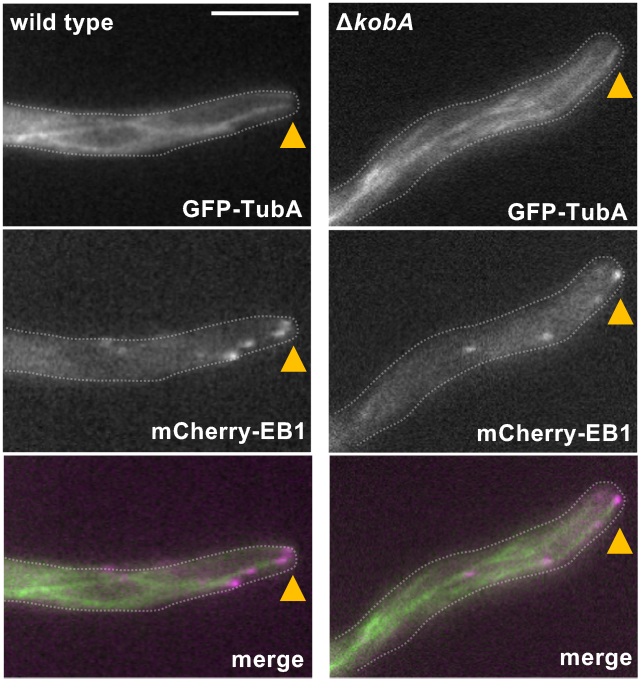
Dual-color imaging of GFP-TubA-marked microtubules and mCherry-EB1-marked microtubule plus ends in a *kobA*+ (wild type) cell and a Δ*kobA* cell. Note that the microtubules in the Δ*kobA* mutant are able to extend to the hyphal tip region. Hyphal tip is indicated by a yellow arrowhead. Bar, 5 μm.

**Figure S4.**
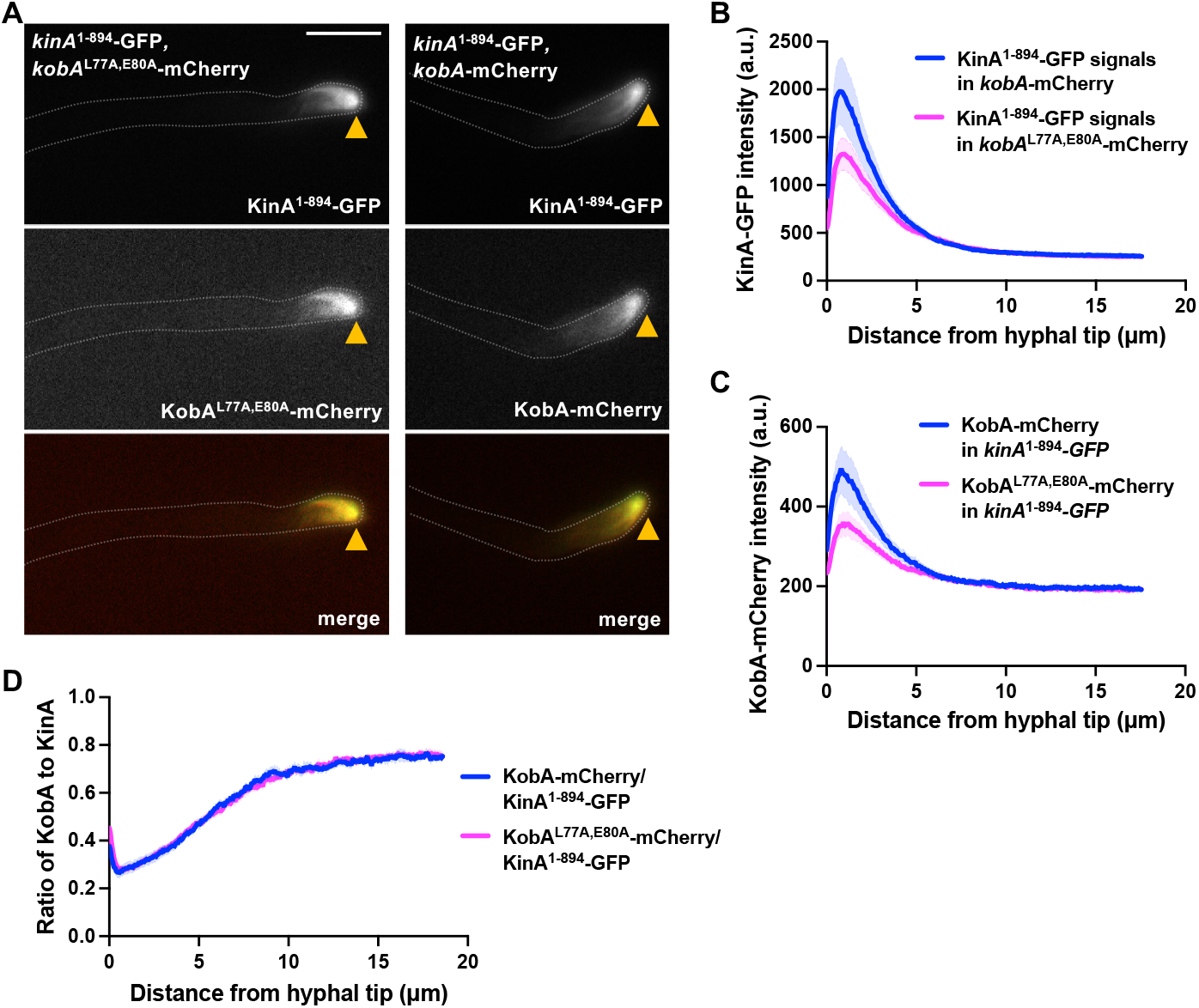
Dual-color imaging and analyses of KinA^1-894^-GFP and KobA-mCherry in the *kobA*^L77A,E80A^-mCherry, *kinA*^1-894^-GFP strain and the *kobA*-mCherry, *kinA*^1-894^-GFP strain. (A) Images of KinA^1-894^-GFP and KobA-mCherry in the *kobA*^L77A,E80A^-mCherry and *kobA*-mCherry backgrounds. Note that the KobA-mCherry signals in *kobA*^L77A,E80A^-mCherry-containing cells are also concentrated at the hyphal tip together with KinA^1-894^-GFP, just like those in the control (*kobA*-mCherry). Hyphal tip is indicated by a yellow arrowhead. Bar, 5 μm. (B) Line scans of fluorescence intensity of KinA^1-894^-GFP in the *kinA*^1-894^*-*GFP, *kobA*-mCherry and the *kinA*^1-894^*-*GFP, *kobA*^L77A,E80A^-mCherry strains. XY graphs with mean (solid lines) and SEM (shading) were generated by Prism 11. Between 0.325 and 0.975 μm from the hyphal tip, KinA^1-894^-GFP intensity was significantly higher in the *kobA*-mCherry strain than that in the *kobA*^L77A,E80A^ strain (p≤0.0005, two-way ANOVA with Šídák’s multiple comparisons test, *n* = 10 hyphae for both strains). (C) Line scans of KobA-mCherry fluorescence intensity in the *kinA*^1-894^*-*GFP, *kobA*-mCherry and the *kinA*^1-894^*-*GFP, *kobA*^L77A,E80A^-mCherry strains. XY graphs with mean (solid lines) and SEM (shading) were generated by Prism 11. Between 0.260 and 1.105 μm from the hyphal tip, KobA-mCherry intensity was significantly higher in the *kobA*-mCherry strain than that in the *kobA*^L77A,E80A^-mCherry strain (p≤0.0005, two-way ANOVA with Šídák’s multiple comparisons test, *n* = 10 hyphae for both strains). (D) Ratios of signal intensity of mCherry to that of GFP along the hypha as a function of distance from the hyphal tip (based on the data used for (B) and (C)). XY graphs with mean (solid lines) and SEM (shading) were generated by Prism 11. There is no significant difference at any of the points analyzed (two-way ANOVA with Šídák’s multiple comparisons test).

**Figure S5.**
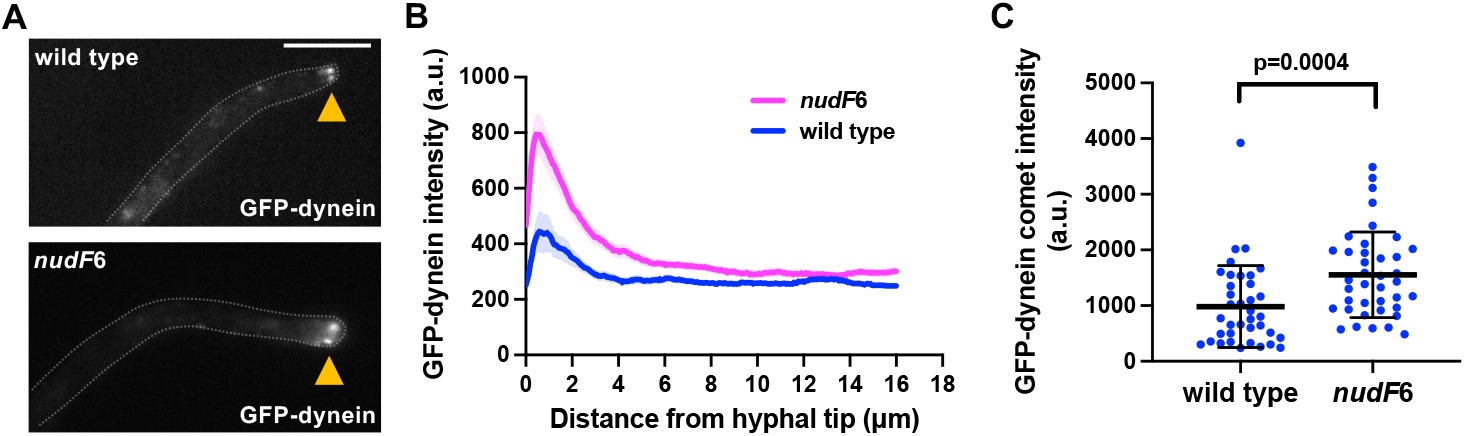
The plus-end dynein accumulation is increased significantly in the *nudF*6 (lis1) mutant. (A) Images of GFP-dynein in wild type and the *nudF*6 background. Hyphal tip is indicated by a yellow arrowhead. Bar, 5 μm. (B) Line scans of fluorescence intensity of GFP-dynein in the wild type and the *nudF*6 backgrounds. XY graphs with mean (solid lines) and SEM (shading) were generated by Prism 11. Between 0 and 2.08 μm from hyphal tip, GFP-dynein intensity was significantly higher in the *nudF*6 strain than that in the wild-type strain (p<0.0001, two-way ANOVA with Šídák’s multiple comparisons test, *n* = 22 hyphae for wild type and *n* = 23 hyphae for *nudF*6). (C) A quantitative analysis of plus-end dynein accumulation (or GFP-dynein comet intensity) in a wild-type strain and a *nudF*6 mutant strain. Scatter plots with mean and SD values were generated by Prism 11, and the p value was generated by Mann-Whitney test (unpaired, two-tailed).

**Figure S6.**
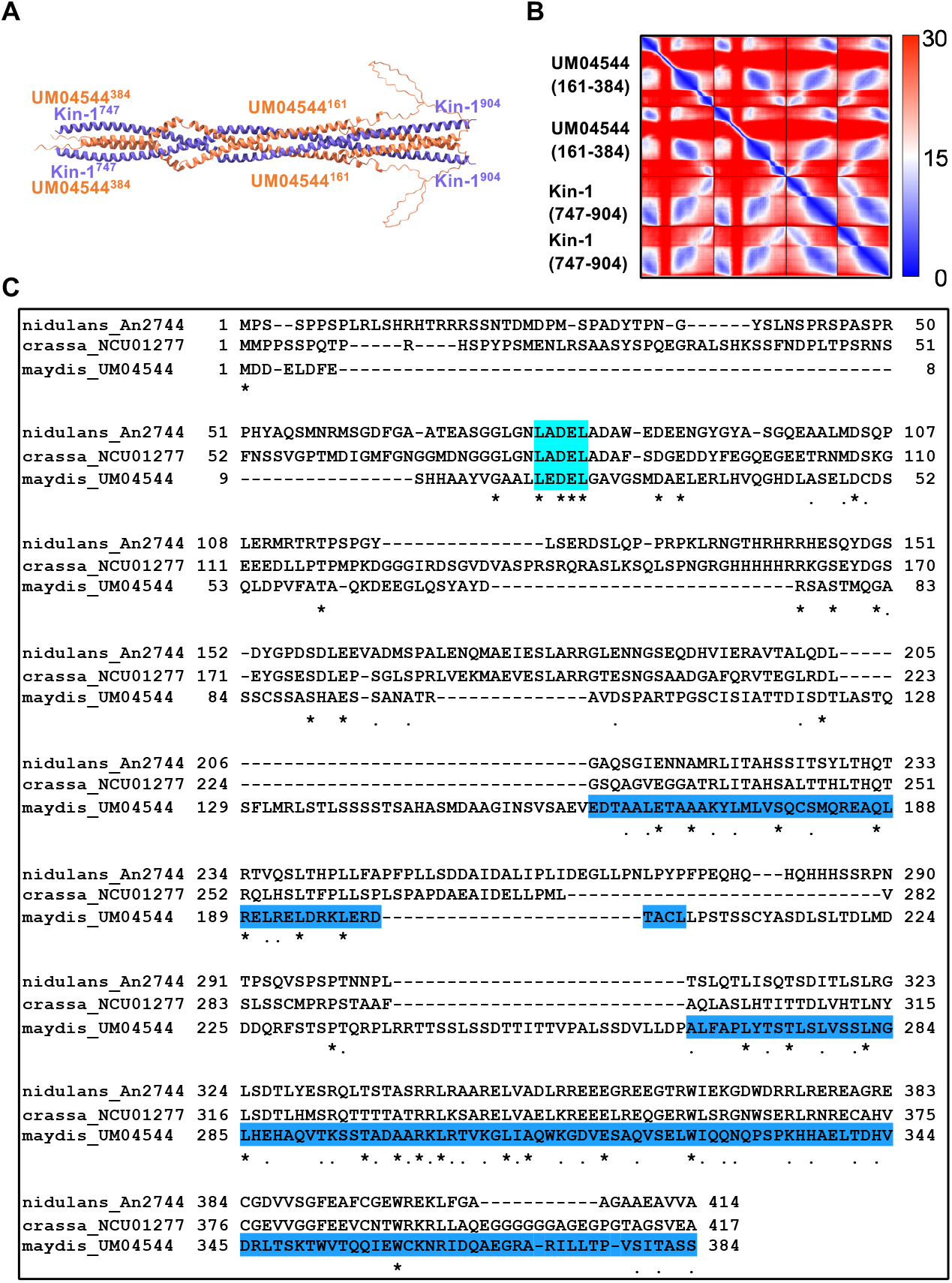
Identification of a *Ustilago maydis* structural homolog of KobA (An2744). (A) AlphaFold-3-based prediction of the interaction between UM04544 and Kin-1 kinesin-1 in *U. maydis*. The C-terminal regions of the UM04544 (aa161-384) dimer and the Kin-1 (aa747-904) dimer interact with each other based on the structural prediction. (B) A predicted aligned error (PAE) plot of the model shown in (A). A lower PAE score, indicated by a darker blue color, indicates a better prediction with lower error. (C) Multiple sequence alignment of KobA (An2744) from *Aspergillus nidulans*, its ortholog from *Neurospora crassa* (NCU01277) and its structural homolog from *U. maydis* (UM04544). The alignment was done using Clustal Omega (1.2.4) at website https://www.ebi.ac.uk/jdispatcher/msa/clustalo. Residues that are identical (*), strongly similar (:) or weakly similar (.) are indicated.

**Table S1.** Mass spectrometry data showing KobA pulled down with KinA in different mutant backgrounds. Numbers of total peptides and unique peptides (unique peptides in parentheses) of proteins pulled down with KinA-GFP (KinA) and KinA^1-894^-GFP (called KinA^K895^* here for simplicity) (a truncation disrupting KinA autoinhibition) ^71^ in different genetic backgrounds such as wild type, Δ*hookA* ^72^, or *alcA*-*nudK*^Arp1^ ^73^, and also those pulled down with ΔC-HookA-GFP (ΔC-HK) ^72^ in wild type and *kinA*^K895^* backgrounds. The results show that KobA can be pulled down with KinA-GFP (KinA) or KinA^1-894^-GFP (KinA^K895^*) and suggest that the KinA-KobA interaction is independent of HookA, dynactin or dynein (based on the near absence of any dynein heavy chain (HC) peptides in the background of *alcA*-*nudK*^Arp1^). In addition, it is important to note that KobA is not pulled down with ΔC-*hookA*-GFP ^72^ (called ΔC-HK here for simplicity) while dynein and dynactin components are pulled down with ΔC-HK. Original data are shown in supplemental excel file 1.

| Protein/ numbers of total<br>and unique peptides | **KinA** | **KinA<sup>K895\*</sup>** | **KinA *∆hookA*** | **KinA<sup>K895\*</sup> *∆hookA*** | **KinA *alcA-nudK<sup>Arp1</sup>*** | **KinA<sup>K895\*</sup> *alcA-nudK<sup>Arp1</sup>*** | **∆C-HK** | **∆C-HK *kinA<sup>K895\*</sup>*** |
|---|---|---|---|---|---|---|---|---|
| **KinA** (An5343, 927aa) | 1732 (91) | 1597 (81) | 1188 (82) | 1608 (77) | 1342 (91) | 1491 (81) | 11 (9) | 13 (10) |
| **KobA (An2744, 414 aa)** | **54 (14)** | **36 (12)** | **29 (10)** | **39 (9)** | **33 (12)** | **36 (12)** | **0 (0)** | **0 (0)** |
| **Dynein HC** (An0118, 4345 aa) | 4 (3) | 23 (18) | 17 (14) | 16 (14) | 0 (0) | 1 (1) | 481 (168) | 381 (154) |
| **Dynein IC** (An1454, 689 aa) | 4 (4) | 5 (5) | 3 (3) | 9 (6) | 5 (5) | 5 (4) | 48 (18) | 46 (18) |
| **Dynactin p50** (An3589, 467 aa) | 3 (3) | 3 (3) | 0 (0) | 4 (4) | 0 (0) | 0 (0) | 52 (18) | 48 (22) |
| **Dynactin p25** (An5022, 202 aa) | 1 (1) | 1 (1) | 1 (1) | 2 (1) | 1 (1) | 1 (1) | 7 (5) | 13 (6) |

## STAR METHODS

### KEY RESOURCES TABLE

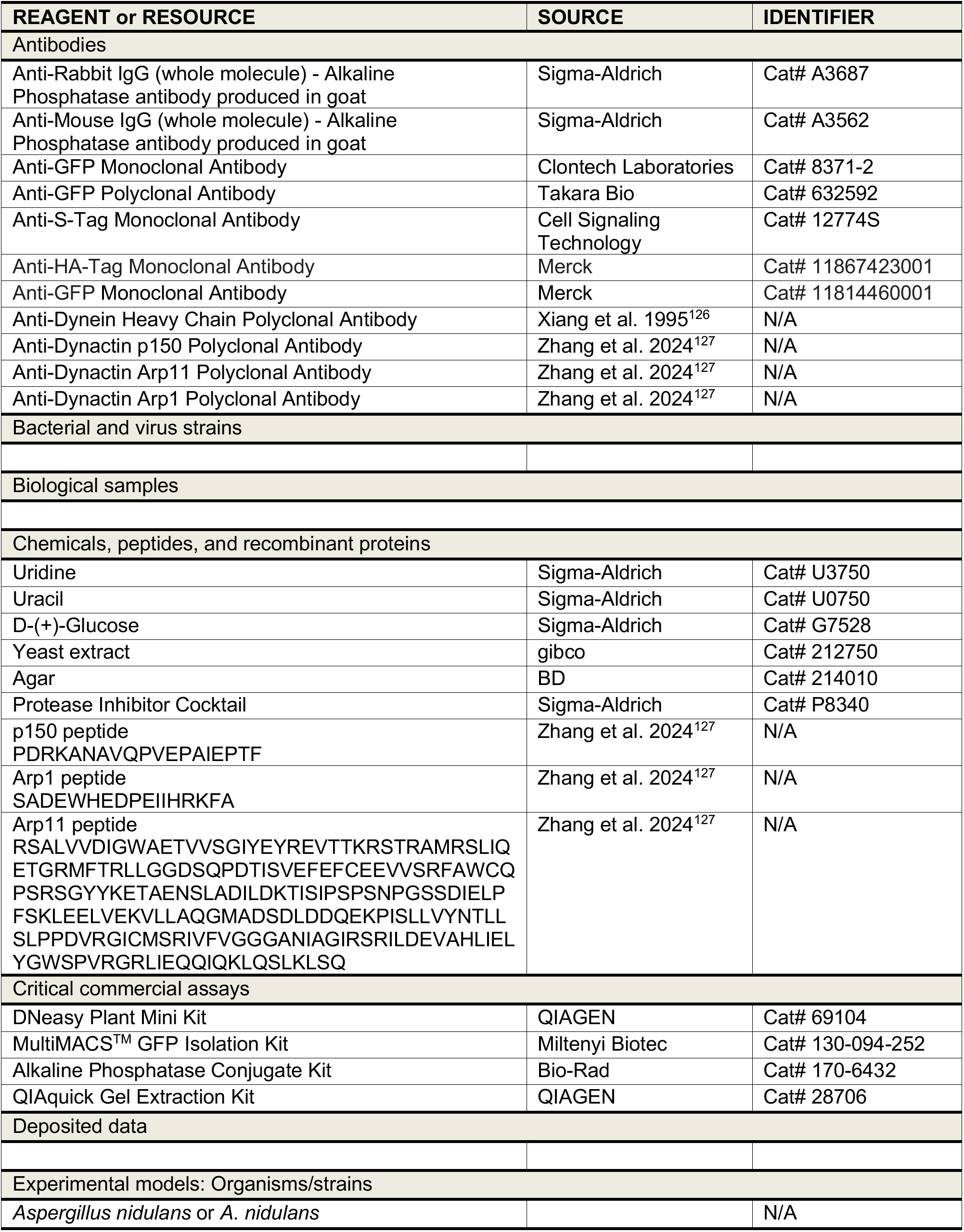
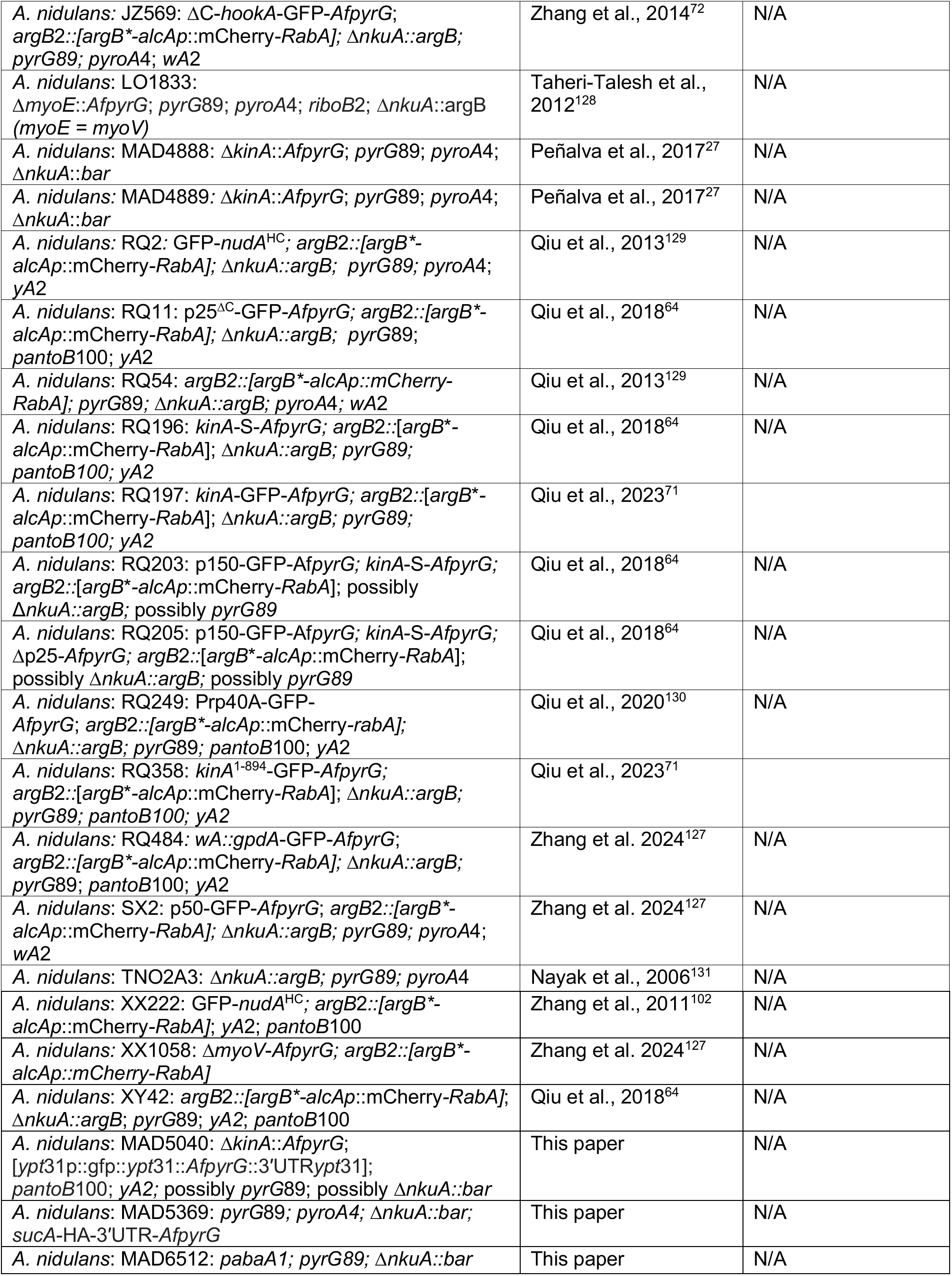
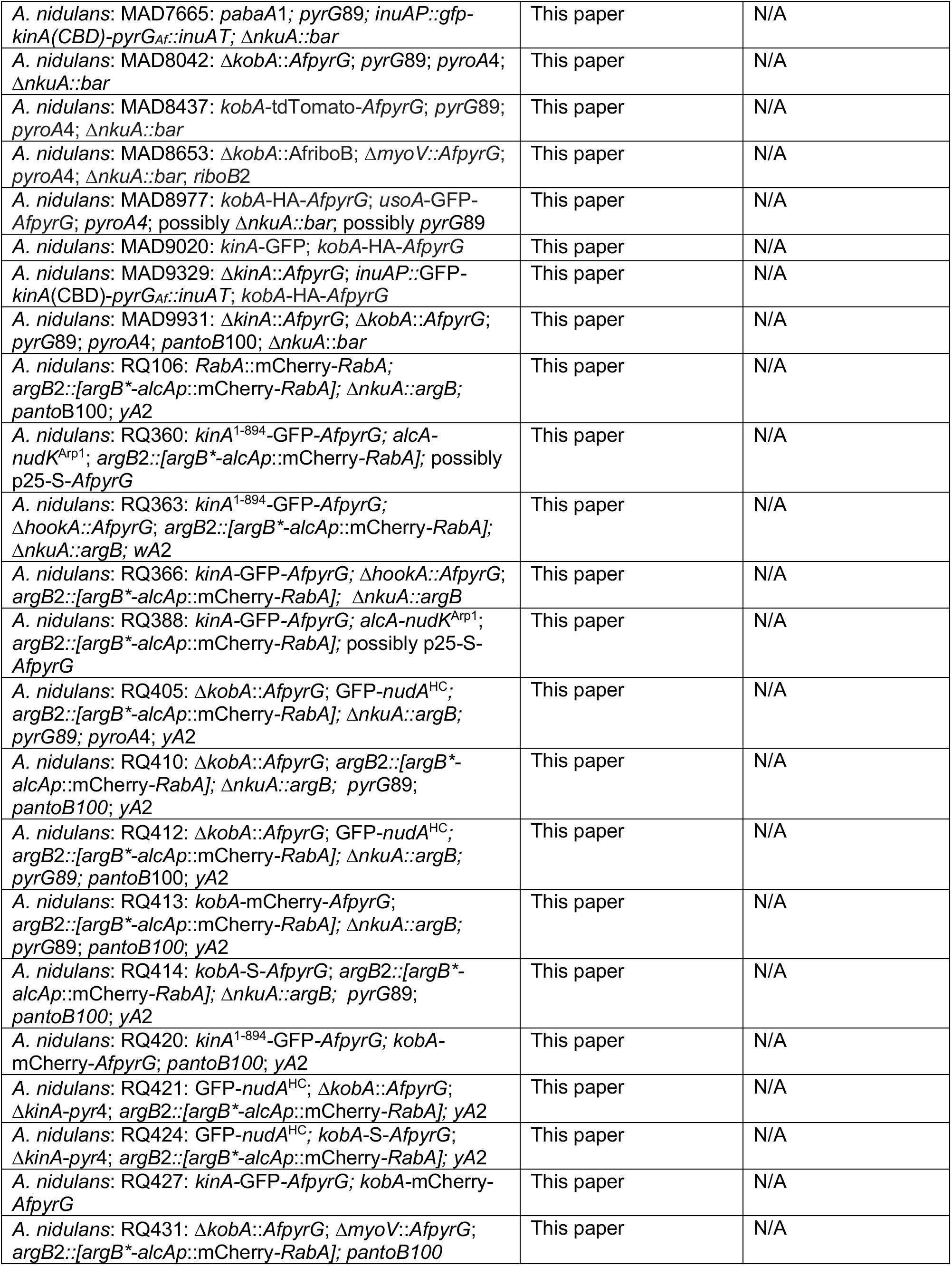
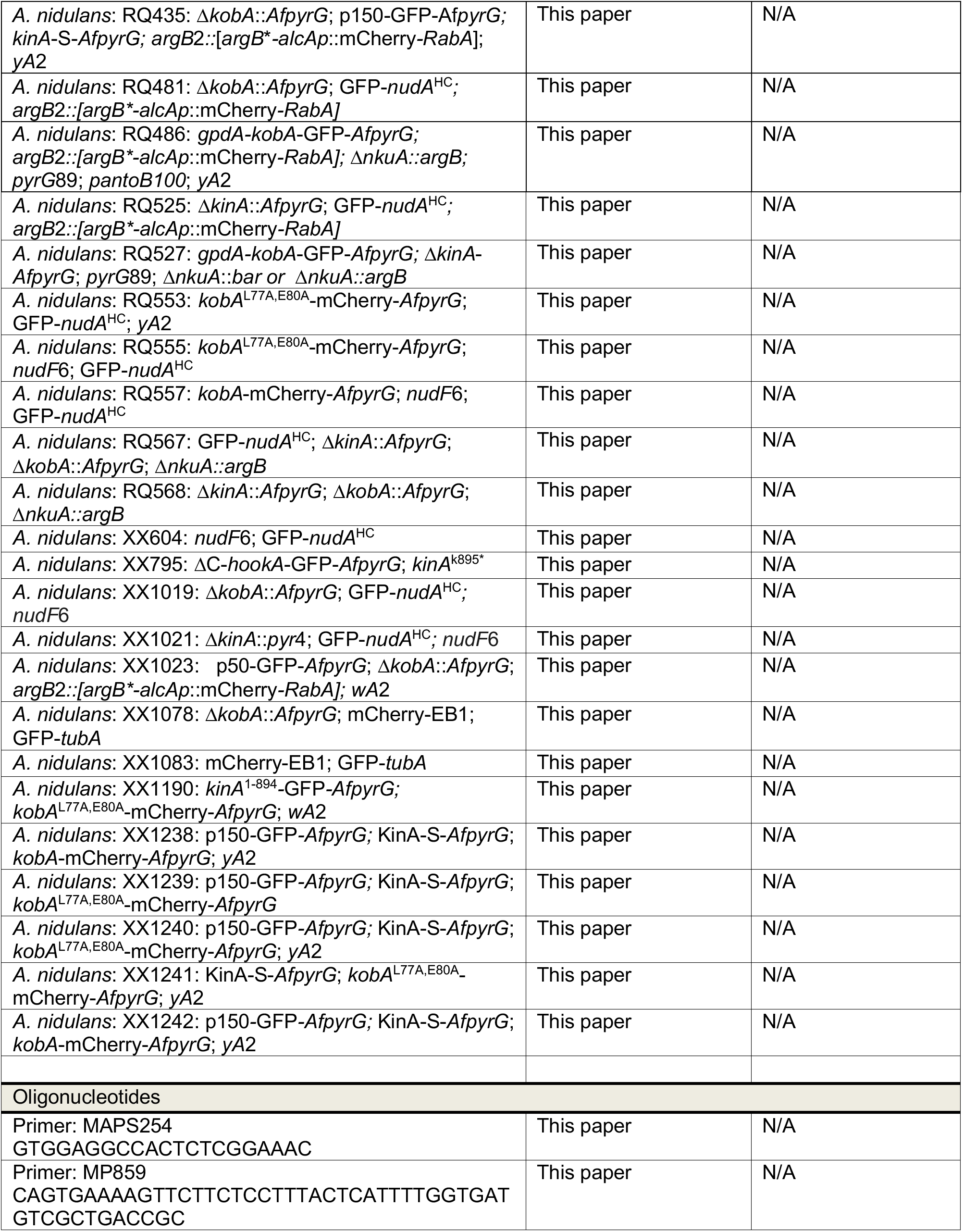
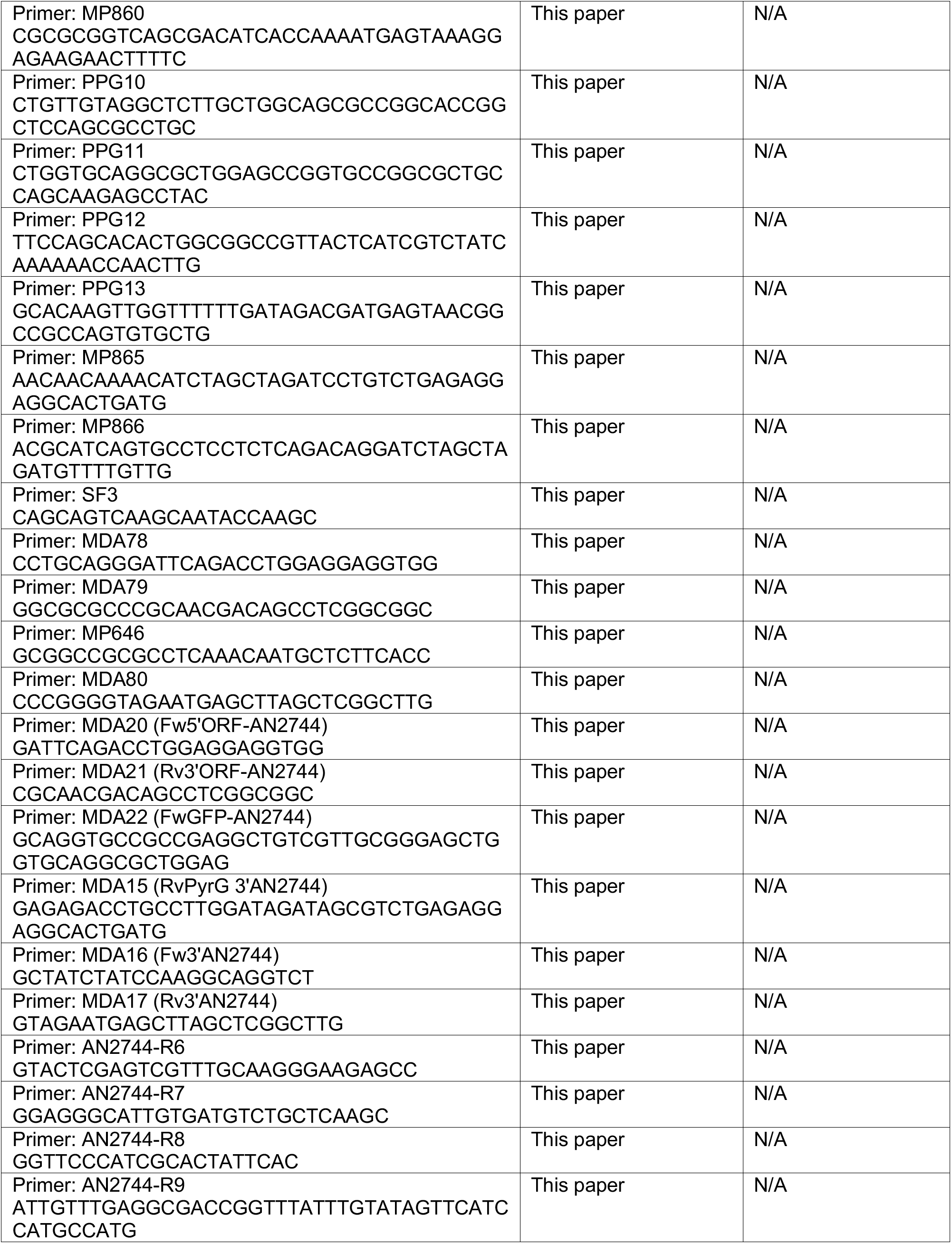
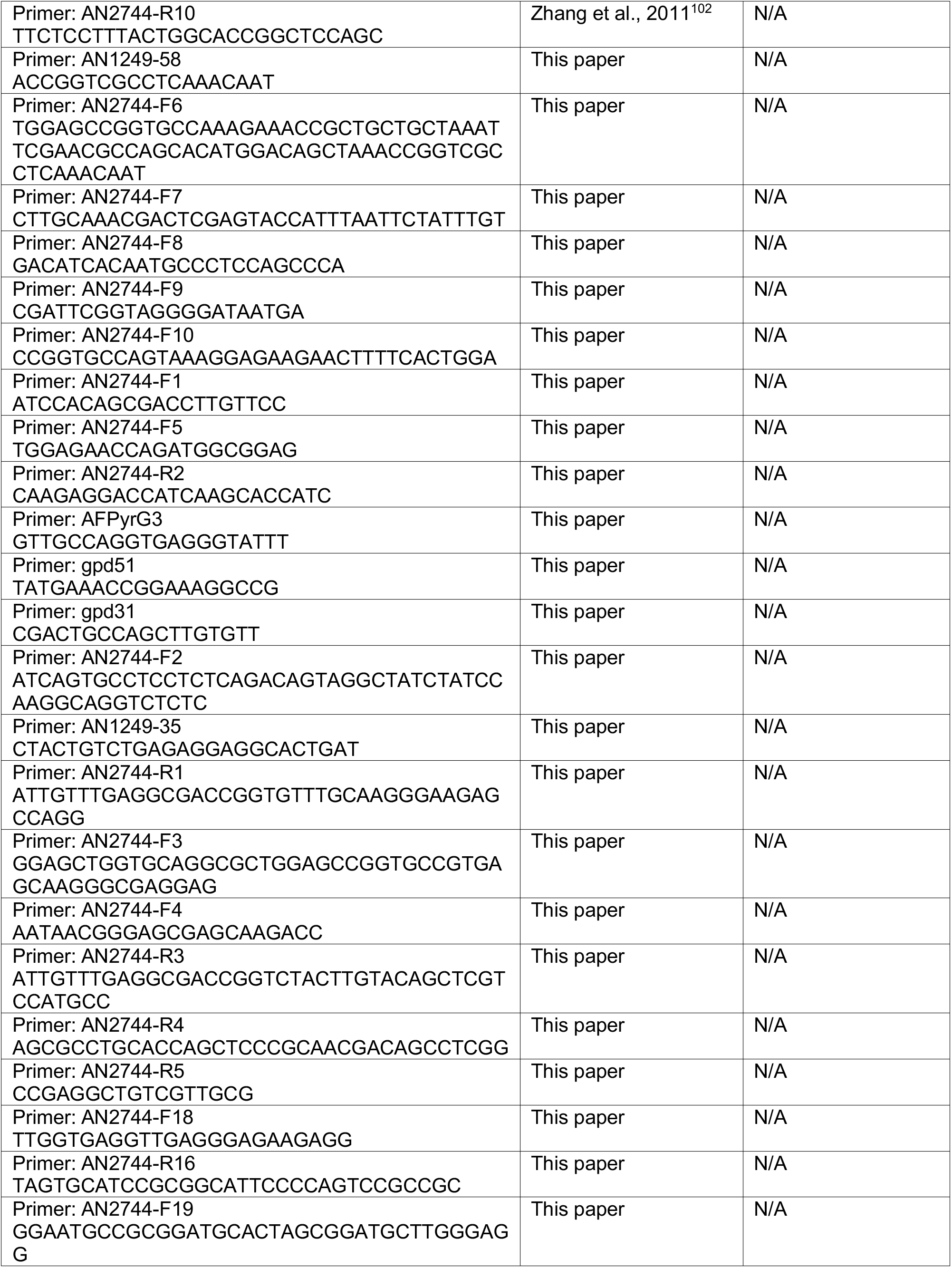
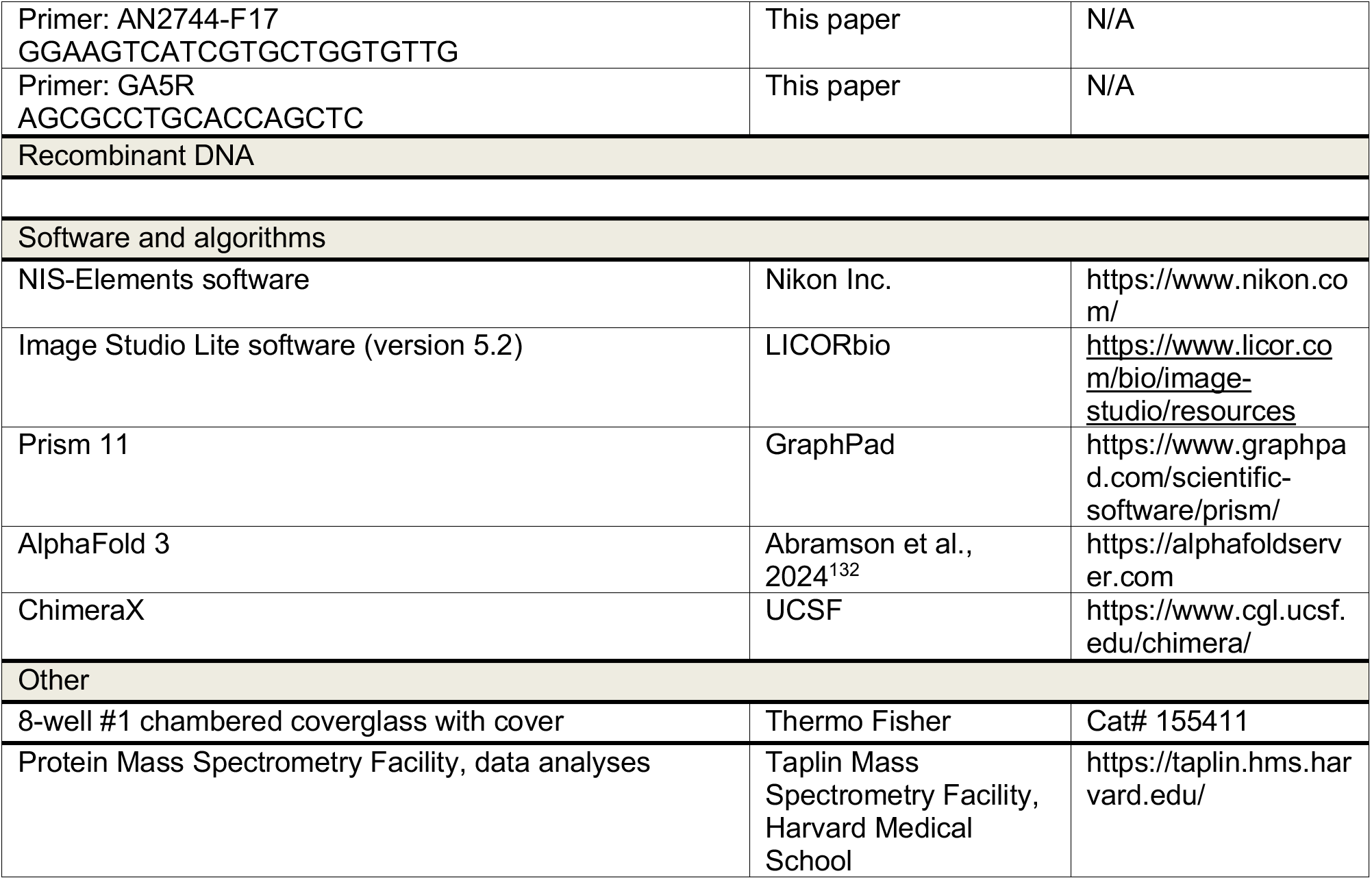

## EXPERIMENTAL MODEL AND STUDY PARTICIPANT DETAILS

The model organism we used for this study is *Aspergillus nidulans*, a filamentous fungus. *A. nidulans* strains grow on solid rich medium made of either YAG (0.5% yeast extract and 2% glucose with 2% agar) or YAG+UU (YAG plus 0.12% uridine and 0.11% uracil) ^133^. Solid minimal medium containing 1% glucose was used for selecting progeny from a genetic cross. Note that the minimal medium (pH 6.5) also contains 0.6% NaNO_3_, 0.052% KCl, 0.0152% KH_2_PO_4_ and 0.051% MgSO_4_. Asexual spores of the strains can be stored in 70% glycerol at - 80°C for several years. For most live-cell imaging experiments, cells were cultured overnight at 32°C in a liquid minimal medium containing 1% glycerol. For the live-cell imaging of *nudF*6-containing strains, we grow the cells at 37°C overnight.

Strains containing multiple mutant alleles were constructed by genetic crosses. Genetic crosses were done by standard methods, and progeny with desired genotypes were selected based on colony phenotype, imaging analysis, western analysis, diagnostic PCR, and/or sequencing of specific regions of the genomic DNA. For genomic DNA isolation, about 0.25 g of hyphal mass was harvested from overnight culture for each sample and pulverized with liquid nitrogen by using mortars and pestles. The DNeasy Plant Mini Kit (QIAGEN) was used for genomic DNA isolation. The QIAquick Gel Extraction Kit (QIAGEN) was used for purifying PCR products from agarose gels. Sequencing was done using the service of QuintaraBio (https://www.quintarabio.com/service/ngs_services).

## METHOD DETAILS

### Live-cell imaging and analyses

Microscopic images used in most figures were generated using a Nikon Ti2-E inverted microscope with Ti2-LAPP motorized TIRF module and a CFI apochromat TIRF 100 x 1.49 N.A. objective lens (oil). The microscope was controlled by NIS-Elements software using 488 nm and 561 nm lines of LUN-F laser engine and ORCA-Fusion BT cameras (Hamamatsu). All the images were taken with a 0.1-s exposure time (binning: 2x2). Image labeling was done using Adobe Photoshop. For all images, cells were grown at 32°C for ∼18 hours in the LabTek Chambered #1.0 borosilicate coverglass system (Nalge Nunc International, Rochester, NY). Images were taken at room temperature, except for experiments using *nudF*6-containing strains in which cells were grown at 37°C and observed using a 37°C chamber. For line scans, we draw a line starting from the hyphal tip in the middle of the hypha and get the average intensity value for the width of 2 μm, which is normally the hyphal width.

### Construction of a strain expressing a GFP-KinA-CBD fusion protein under the control of the inuAp

A linear DNA molecule consisting, from 5’ to 3’, of the promoter region of the inuA gene, the sGFP coding region, codons 731-927 of KinA fused in frame to the C-terminus of sGFP, the *pyrG* gene of *A. fumigatus* (*pyrGAf*) and the *inuA* 3’ UTR, was assembled by 5-way fusion PCR and used to replace the resident *inuA* locus by homologous recombination. The promoter region including 857 bps upstream of the InuA ATG codon was PCR-amplified with primers MAPS254 and MP859; the sGFP coding region was amplified with primers MP860 and PPG10; the sequence encoding the CBD of KinA (644 bp, amino acids 731-927) was amplified from cDNA from a wild-type strain using the oligo pair PPG11 and PPG12; the pyrG gene from A. fumigatus (1949 bp) was amplified with oligos PPG13 and MP865; and the 851 bp *inuA* 3’ UTR was amplified with primers MP866 and SF3. Fragments were mixed in appropriate proportions, and a 5.8 kb cassette was assembled using primers MAPS254 and SF3, which was used to transform MAD6512; transformants were genotyped by diagnostic PCR, and strain MAD7665 was selected for subsequent experiments.

### Endogenous tagging of KobA with HA3

A strain expressing endogenously HA3-tagged KobA was constructed by homologous recombination–mediated gene replacement using a cassette for C-terminal HA3 tagging, which contained the *pyrG* gene from *A. fumigatus*. The cassette was generated by 3-way fusion PCR (oligo pairs MDA20 and MDA21; MDA22 and MDA15; and MDA16 and MDA17) and used to transform a recipient Δ*nkuA* strain (deficient in the non-homologous end-joining pathway)^131^. Correct integration was confirmed by PCR using external primers, and the strain with the correct integration was chosen for subsequent work.

### Tagging of KobA (An2744) with tdTomato

We used a template plasmid in which the coding region for a (Gly-Ala)5-tdTomato fusion polypeptide was flanked by 5′SbfI-AscI and 3′NotI-XmaI restriction sites^134^. The 3’ region of kobA (An2744) without the stop codon (amplified with primers MDA78 and MDA79 containing SbfI and AscI restriction sites respectively) was cloned into the (Gly-Ala)5-tdTomato plasmid, generating an intermediate plasmid. The pyrGAf gene fused with the 3′ UTR region of *kobA* was amplified with primers MP646 and MDA80 containing NotI and XmaI restriction sites, respectively. This fragment was cloned in the intermediate plasmid after NotI and XmaI digestion. The complete cassette was excised after digestion with SbfI and XmaI and used to transform a Δ*nkuA* strain.

### Construction of a Δ*kobA* (ΔAn2744) strain

The ΔAn2744-afpyrG replacement cassette was generated by fusion PCR using six oligonucleotides: AN2744-F1, AN2744-F2, AN1249-58, AN1249-35, AN2744-R1, and AN2744-R2. Genomic DNA from strain RQ249, which contains an AN1249-GFP fusion, was used as template. AN2744-F1 and AN2744-R1 amplified a ∼1.0 kb fragment corresponding to the 5’ UTR of An2744. AN1249-58 and AN1249-35 amplified a ∼1.9 kb fragment linking the 5’ UTR to *afpyrG*, and AN2744-F2 and AN2744-R2 amplified a ∼0.7 kb fragment connecting *afpyrG* to the 3’ UTR. Fusion PCR with primers AN2744-F1 and AN2744-R2 generated a ∼3.6 kb ΔAn2744-afpyrG cassette, which was used to transform strain RQ2 (GFP-nudA).

Transformants were screened by fluorescence microscopy for GFP-nudA signal. Correct gene replacement at the *An2744* locus was confirmed by diagnostic PCR using primers AN2744-F1 and AN2744-R2, which produced an amplicon of ∼3.6 kb in the ΔAn2744 strain compared with a ∼3.0 kb fragment in the wild-type strain.

### Construction of a strain carrying *kobA*(An2744)-S at the *kobA* locus

The An2744-S-afpyrG cassette was generated by fusion PCR using the following oligonucleotides: AN2744-F6, AN2744-R5, AN2744-F4, AN2744-R2. AN2744-F4 and AN2744-R5 were used to amplify a ∼0.8 kb fragment from wild-type genomic DNA. AN2744-F6 and AN2744-R2 were used to amplify a ∼2.7 kb fragment from genomic DNA of a Δ*kobA* strain containing the *AfpyrG* gene. AN2744-F4 and AN2744-R2 were used for fusion PCR to fuse the two fragments and generate a ∼3.4 kb fragment, which contains the sequence encoding the 5 x glycine and alanine repeats followed by an S-tag before the *kobA* stop codon. This fragment was used to transform strain XY42.

Correct integration at the An2744 locus was confirmed by diagnostic PCR using primers AN2744-F5, and AFPyrG3, which yielded an amplicon of ∼0.9 kb, and by sequencing of this fragment using AN2744-F5 and AFPyrG3.

### Construction of a strain carrying the *gpdA*-*kobA*(An2744)-GFP at the *kobA* locus

The gpdA-An2744-GFP-afpyrG cassette was generated by fusion PCR using ten oligonucleotides: AN2744-R6, AN2744-R7, AN2744-R8, AN2744-R9, AN2744-R10, AN1249-58, AN2744-F7, AN2744-F8, AN2744-F9 and AN2744-F10. Genomic DNA from strain RQ414 (An2744-S) served as template. AN2744-F9 and AN2744-R6 amplified a ∼0.8 kb fragment corresponding to the 5’ untranslated region (UTR), AN2744-F7 and AN2744-R7 amplified a ∼1.2 kb fragment containing the *gpdA* promoter, AN2744-F8 and AN2744-R10 amplified a ∼1.4 kb fragment of the An2744 coding region, and AN1249-58 and AN2744-R8 amplified a ∼2.7 kb fragment containing *afpyrG* fused to the 3’ UTR. AN2744-F10 and AN2744-R9 amplified a ∼0.7 kb GFP fragment from genomic DNA of strain RQ11 (p25^ΔC^-GFP). The five fragments were fused by fusion PCR using primers AN2744-F9 and AN2744-R2 to generate a ∼6.6 kb gpdA-An2744-GFP-afpyrG fragment, which was used to transform strain XY42.

Transformants were pre-screened by fluorescence microscopy for GFP signal. Correct integration at the An2744 locus was confirmed by diagnostic PCR using primers AN2744-F1 and AN2744-R8, which yielded an amplicon of ∼6.9 kb, and by sequencing of this fragment. Absence of additional mutations within the cassette was verified using sequencing primers AN2744-F5, AFPyrG3, gpd51 and gpd31.

### Construction of a strain carrying KobA-mCherry (An2744-mCherry) at the *kobA* locus

The An2744-mCherry-afpyrG cassette was assembled by fusion PCR using primers AN2744-F3, AN2744-F4, AN1249-58, AN2744-R2, AN2744-R3 and AN2744-R4. Genomic DNA from strain RQ106 (mCherry-RabA) was used as template for tagging components. AN2744-F3 and AN2744-R3 amplified a ∼0.8 kb fragment of mCherry, and AN2744-F4 and AN2744-R4 amplified a ∼0.8 kb fragment corresponding to the C terminus of the *An2744* coding region. AN1249-58 and AN2744-R2 amplified a ∼2.6 kb fragment containing *afpyrG* linked to the 3’ UTR from strain RQ405 (ΔAn2744). Fusion PCR using primers AN2744-F4 and AN2744-R2 yielded a ∼4.1 kb An2744-mCherry-afpyrG cassette, which was used to transform strain XY42. Transformants were screened for mCherry fluorescence, and correct integration at the *An2744* locus was verified by diagnostic PCR using primers AN2744-F5 and AFPyrG3.

### Construction of a strain carrying KobA(An2744)^L77A,E80A^-mCherry at the *kobA* locus

The An2744^L77A,E80A^-mCherry-afpyrG cassette was generated by fusion PCR using genomic DNA from strain RQ427 (An2744-mCherry) as template. Primers AN2744-F18 and AN2744-R16 amplified a ∼0.8 kb fragment containing the 5’ UTR and the N-terminal region of An2744, while AN2744-F19 and AN2744-R2 amplified a ∼4.4 kb fragment comprising the remaining portion of the An2744 coding region (carrying the L77A and E80A mutations), mCherry, afpyrG, and the 3’ UTR. Fusion PCR with primers AN2744-F17(5’-GGAAGTCATCGTGCTGGTGTTG-3’) and AN2744-R2 produced a ∼5.2 kb An2744^L77A,E80A^-mCherry-afpyrG fragment, which was used to transform strain XY42.

Transformants were screened for mCherry fluorescence. Correct integration at the *An2744* locus was confirmed by diagnostic PCR using primers AN2744-F18 and AFPyrG3, yielding a ∼2.7 kb fragment. The amplified product was sequenced with primers AN2744-F18 and GA5R to verify introduction of the L77A and E80A substitutions and to exclude additional mutations.

### Biochemical pulldown assays, western analysis and antibodies

To harvest cells, overnight grown fungal mycelia (or hyphal mass) were filtered through miracloth (EMD Millipore Corp.) and washed with ∼200 ml distilled water. Excess liquid was removed by pressing the folded miracloth (containing the hyphal mass inside) between paper towels. After the liquid has been removed, the weight of the hyphal mass (without the miracloth) was measured on a balance, and about 0.6 g hyphal mass was collected and wrapped with a foil paper, labeled and stored in a -80°C freezer. To break hyphal cells, a mortar and pestle were used to grind the frozen 0.6 g hyphal mass to a fine powder in liquid nitrogen (basically, liquid nitrogen was added in small amounts during grinding to keep the sample frozen). Cell extracts were prepared by adding 1.5 mL of lysis buffer containing 50 mM Tris-HCl (pH 8.0), 0.01% Triton X-100 and 10 μg/mL of a protease inhibitor cocktail (Sigma-Aldrich). Cell extracts were centrifuged at 20,000 *g* for 60 minutes at 4°C, and supernatant was used for the pull-down experiment. The MultiMACS^TM^ GFP Isolation Kit (Miltenyi Biotec) was used to pull down proteins associated with the GFP-tagged protein, which was done as described previously ^72^. To pull down GFP-tagged proteins, 35 μL anti-GFP MicroBeads were added into the cell extracts for each sample and incubated (with gentle rotating) at 4°C for 60 minutes. The MicroBeads/cell extracts mixture was then applied to the μColumn followed by gentle wash with 200 μL of the same lysis buffer used above for protein extraction. After washing three times, 80 μL of 2 x SDS-PAGE sample buffer was used to elute the proteins with boiling. The eluted proteins were applied to premade 4–15% Criterion™ TGX Stain-Free™ Protein Gels (Bio-Rad), and after electrophoresis, proteins in the gels were transferred overnight at 4°C to nitrocellulose membrane (0.45 μm) (Bio-Rad). Western blot analyses were performed using the alkaline phosphatase system and blots were developed using the AP color development reagents from Bio-Rad. Quantitation of the protein band intensity was done using the Image Studio Lite software (version 5.2). Specifically, an area containing the whole band was selected as a region of interest, and the intensity sum within the region of interest was measured. Then, the region of interest box was dragged to the equivalent region of the negative control lane or a blank region without any band on the same blot to take the background value, which was then subtracted from the intensity sum.

The rabbit polyclonal antibody against GFP was from Takara Bio Inc (Catalog number: 632592). The rabbit monoclonal antibody against the S-tag was from Cell Signaling Technology (Catalog number: 12774S). Polyclonal antibody against Arp1 was generated using the service of Pacific Immunology (www.pacificimmunology.com). An immunograde peptide of Cys-SADEWHEDPEIIHRKFA from the C-terminus of Arp1 was synthesized and conjugated to the KLH carrier protein, and this conjugated form was used as an antigen for rabbit injection. The antibody was affinity-purified using the same peptide. Similarly, polyclonal antibody against p150 was generated using the service of Pacific Immunology (www.pacificimmunology.com), and an immunograde peptide of Cys-PDRKANAVQPVEPAIEPTF from the C-terminus of p150 was used. Polyclonal antibody against Arp11 was generated using the service of Boster Bio (www.bosterbio.com). A recombinant protein of 217 amino acids (RSALVVDIGWAETVVSGIYEYREVTTKRSTRAMRSLIQETGRMFTRLLGGDSQPDTISVEFEFC EEVVSRFAWCQPSRSGYYKETAENSLADILDKTISIPSPSNPGSSDIELPFSKLEELVEKVLLAQ GMADSDLDDQEKPISLLVYNTLLSLPPDVRGICMSRIVFVGGGANIAGIRSRILDEVAHLIELYGW SPVRGRLIEQQIQKLQSLKLSQ) was used for rabbit injection and purification of serum.

The methods for pulldown for the experiments with GFP-KinA-CBD are as follows: Lyophilized mycelium (70 mg) was disrupted in 2 mL tubes containing a ceramic bead using a FastPrep instrument (15 s pulse, power 4) until a fine powder was obtained. This powder was resuspended in 1.5 mL of extraction buffer (25 mM Tris-HCl pH 7.5, 150 mM NaCl, 0.5 mM EDTA, 0.5% NP-40) supplemented with cOmplete™ ULTRA protease inhibitor cocktail tablets (Sigma-Aldrich, #05892953001), Mg132 (5 μM), and ∼100 μL of 0.6 mm glass beads. The samples were homogenized using a FastPrep (15 s pulse, power 6). The homogenate was incubated (10 min at 4°C) on a rotating wheel. This extraction step was repeated two more times. The resulting homogenate was clarified by centrifugation (15,000 × g for 30 min at 4°C) using a refrigerated microcentrifuge. Approximately 100 mg of total protein (corresponding to ∼5 mL of extract) was incubated with 50 µL of GFP-Trap magnetic agarose beads (Chromotek, #gtma-20) for 2 hours at 4°C on a rotating wheel. The beads were washed three times with washing buffer (25 mM Tris-HCl pH 7.5, 150 mM NaCl, 0.5 mM EDTA) using a magnet and diluted in the same buffer. Magnetic GFP-Trap agarose beads were separated using a magnet. This separation step was repeated two more times using 1 mL of washing buffer. The final pellet was resuspended in 60 µL of lysis buffer. The sample was boiled for 5 min and centrifuged to remove the GFP-Trap magnetic agarose beads. Finally, 15 µL of each sample were resolved by SDS-PAGE, transferred to a membrane and analyzed by Western blot using anti-HA (Merck) and anti-GFP (Merck) antibodies.

### Statistical analysis

All statistical analyses were done using GraphPad Prism 11 for macOS. The D’Agostino & Pearson normality test was performed on all data sets except for data sets with small n (n=3). For western blot data, data distribution was assumed to be normal but this was not formally tested because the n number is 3 for these data sets. For these data, Welch’s t test (unpaired, two-tailed) was used for analyzing any two data sets, and Ordinary one-way ANOVA with Dunnett’s multiple comparison test was used for analyzing multiple data sets. For all other data sets, the D’Agostino & Pearson normality test was performed. The data sets shown in Figure 6D and Figure S5C did not pass the D’Agostino & Pearson normality test (alpha=0.05) and were analyzed using the Mann-Whitney test (two-tailed), a non-parametric test without assuming Gaussian distribution. The data sets shown in Figure S1 did not pass the D’Agostino & Pearson normality test (alpha=0.05) and were analyzed using the Kruskal-Wallis ANOVA test (with Dunn’s multiple comparisons test, unpaired), a non-parametric test without assuming Gaussian distribution. For analyzing the line scans of two groups, we used two-way ANOVA with Šídák’s multiple comparisons test to compare each cell mean with other cell mean in that row (Note that each row represents a position with a particular distance from the hyphal tip). For analyzing the line scans of multiple groups, we used two-way ANOVA with Tukey’s multiple comparisons test to compare each cell mean with the other cell mean in that row.

### AlphaFold-3-based structural prediction

Protein three-dimensional structures were predicted using AlphaFold-3 ^132^ (accessed via the AlphaFold Server, https://alphafoldserver.com) in template-free mode. Predicted models were subsequently inspected and analyzed using UCSF ChimeraX.

